# Characterization of the functional and clinical impacts of CACNA1A missense variants found in neurodevelopmental disorders

**DOI:** 10.1101/2025.04.23.650298

**Authors:** Erkin Kurganov, Lei Cui, Nikita Budnik, Siwei Chen, Erick Olivares, David Baez-Nieto, Laina Lusk, Lacey Smith, Sooyeon Jo, Sureni V. Mullegama, Amanda Lindy, Alfred L George, Annapurna Poduri, Ingo Helbig, Mark Daly, Jen Q. Pan

## Abstract

*CACNA1A* encodes the P/Q-type Ca_V_2.1 calcium channels whose function underlies neuronal excitability, presynaptic neurotransmitter release, and Ca^2+^ signaling in neurons. Pathogenic variants in *CACNA1A* have been found in individuals with various neurological conditions, including hemiplegic migraine, epilepsy, developmental delay, and ataxia. Clinical presentations can vary significantly between patients, with limited information known about the underlying neurobiology of these different clinical patterns. Adding further complication, prior work on pathogenic missense variants has demonstrated variable impacts on CaV2.1 channel function, sometimes in opposite directions. As such, the relationships between specific coding variants, electrophysiological properties, and clinical phenotypes remain elusive. In this study, we determined the biophysical properties of an allelic series of 42 *de novo* missense *CACNA1A* variants discovered in a neurodevelopmental disorder cohort of more than 31,000 individuals, together with the most common eight coding variants found in the general population. We found that all but one *de novo* variant altered at least one aspect of the channel properties examined, and the majority (70%) of the variants reduced the channel current density. In addition, for variants that encode human Ca_V_2.1 channels (hCa_V_2.1) with detectable currents, nearly 50% altered how channels respond to membrane potential, while common variations did not significantly change any channel biophysical properties. Coupled with our functional analyses and AlphaMissense prediction, we showed that Ca_V_2.1 missense variants significantly underlie the risk of developmental epileptic encephalopathy. Subsequently, we examined the physiological impact of variant hCa_V_2.1 using NEURON simulations as an omnibus output of neuronal function and found that abnormal biophysical channel properties have a profound impact on Purkinje cell excitability. Most interestingly, we correlated the clinical phenotype with molecular consequences of missense variants provided by our comprehensive functional analyses and found that distinct Ca_V_2.1 channel molecular function is significantly associated with different clinical outcomes. By analyzing an entire allelic series of *CACNA1A de novo* changes in a large cohort of individuals with neurodevelopmental disorders, we provide a powerful approach to dissecting the role of missense variants in *CACNA1A* channelopathy, which in turn may help pave the way for future precision medicine initiatives.

## INTRODUCTION

*CACNA1A* encodes the P/Q-type Ca_V_2.1 calcium channels expressed throughout the nervous system. Ca_V_2.1 channels serve as the major calcium influx mechanism that underlies neuronal excitability, presynaptic neurotransmitter release, and postsynaptic Ca^2+^ signaling in dendrites and cell bodies of many neurons(*1–3*). As such, they play a critical role in normal neuronal development and diseases. Coding sequence mutations of *CACNA1A* have been found in patients with familial hemiplegic migraine (FHM1)(*4–8*), episodic ataxia (EA2)(*5, 9*), and spinocerebellar ataxia (SCA6)(*9, 10*). For example, recurrent *CACNA1A* missense mutations that alter voltage dependence of activation(*11, 12*) and inactivation kinetics(*13, 14*) were found in FHM1 (OMIM: 141500), whereas protein truncating mutations have been associated with EA2 (OMIM: 108500)(*15*). More recently, both gain-of-function and loss-of-function *de novo* missense *CACNA1A* variants were associated with Lennox-Gastaut syndrome(*16*), a rare but severe type of epilepsy. As missense coding changes of *CACNA1A* may disrupt channel function with distinct and sometimes opposite effects, it is unclear whether different molecular phenotypes of Ca_V_2.1 function underlie similar, distinct, or overlapping clinical phenotypes.

In this study, we determined the functional impact of an allelic series of 36 *de novo* missense variants of *CACNA1A* found in a large cohort of 31,058 parent-offspring trios of individuals with severe developmental disorders. Each proband was heterozygous for *a de novo* coding variant in *CACNA1A* (*17*). In addition, we included 6 novel *de novo CACNA1A* variants from pediatric patients in the functional analyses. Finally, we included 8 missense variants from the gnomAD database with minor allele frequencies greater than 0.2% to serve as control Ca_V_2.1 channel properties in the general population. To study the functional impact of missense coding variants of *CACNA1A*, we expressed and analyzed essential biophysical properties of each variant-encoded Ca_V_2.1 channel using whole cell voltage clamp recording (*18*) and compared to those of the wild-type reference channels. We identified which and to what extent the biophysical properties of the Ca_V_2.1 channel were impacted by common and *de novo* disease-associated missense variants. The biophysical properties of Ca_V_2.1 variant channels were then used to model the impact of the missense variants on simulated Purkinje neuron excitability(*19*) as a measure of physiological output. Furthermore, using AlphaMissense, a newly developed computational algorithm to predict the pathogenicity of missense variants, we evaluated the overall contribution of *CACNA1A* missense variants to severe neurodevelopmental disorder risk. Most notably, we obtained clinical information from patients who carry some of the variants that we characterized and sought to clarify the relationship between molecular function of Ca_V_2.1 with clinical phenotype. Such functional analyses of an entire allelic series of *de novo* missense variants of Ca_V_2.1 channels, in the context of human genetics, strongly support that pathogenic missense coding changes of *CACNA1A* significantly contribute to the genetic risk of neurodevelopmental disorders. These analyses associate specific molecular function of Ca_V_2.1 to certain clinical phenotypes, provide important insight into the role of *CACNA1A* in disease risk, and pave the way for precision treatment for patients who carry *CACNA1A* mutations.

## RESULTS

### De novo missense variants and common variants of CACNA1A reside in distinct domains of the channel protein

We analyzed 50 missense variants of *CACNA1A*, of which 42 are disease-associated and 8 are common in the general population. Among 42 missense variants we characterized, 36 missense changes were identified in 55 patients in a large cohort of neurodevelopmental disorders, representing an entire missense allelic series of *de novo* coding changes in *CACNA1A* within 31,058 parent–offspring trios(*17*). Six missense variants (V176M, A713T, R1349Q, V1393M, D1634N, and R1664Q) were *de novo* and recurrent in this cohort. In addition, six novel *de novo* variants were identified in patients with neurodevelopmental disorders outside of this cohort, whose impact on channel function was unknown and we included them in our analyses (see method). To estimate the functional range of properties of the Ca_V_2.1 channel in the general population, we included the most frequent common coding variants of *CACNA1A* annotated in the gnomAD database (Table 1), with minor allele frequencies exceeding 0.2%. The most common coding change of *CACNA1A* was E992V, comprising 13.5% of the total alleles in the general population.

**Table I.**
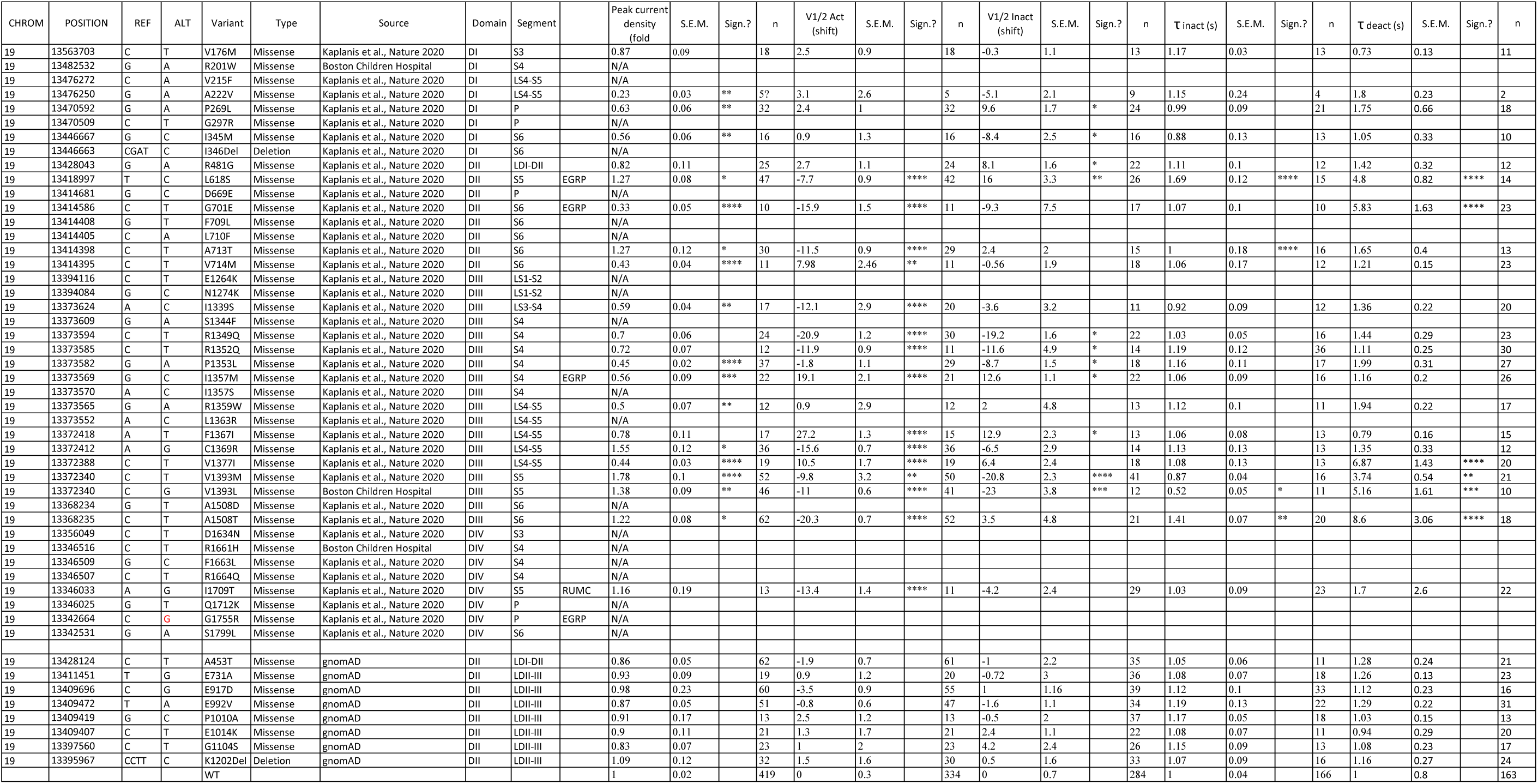
Functional consequences of de novo variants of CACNA1A. The genomic location, allele frequencies and structure (domain and segment) locations of CACNA1A variants are shown. The average functional impact of each variant of the hCav2.1 channels compared to WT channels is shown for each biophysical property (mean ± S.E.M.). Biophysical properties were left blank for of variants that did not produce currents. “*” indicates significant difference from WT channels using One-way ANOVA, Dunnett’s post hoc <0.05 for each variant in each biophysical property. CHTOM = Chromosome POSITION = Position within the chromosome REF = Reference nucleotide ALT = Nucleotide change Variant = amino acid change Source = Clinical classification Domain = One of the four domains of hCav2.1 channel Segment = One of the six segments of the domain Peak current = Normalized peak current change respect to WT V1/2 Act = Voltage dependence of activation shift respect to WT V1/2 Inact = Voltage dependence of inactivation respect to WT τ inact = Tau inactivation τ deact = Tau deactivation S.E.M. = Standard mean error Sign.? = Statistical significance n = total number of cells analyzed N/A = not available (currents for these variants were too little to be reliably measured)

*CACNA1A* encodes the core α1 subunit of the human Ca_V_2.1 (hCa_V_2.1) channel, comprised of four pseudo symmetric transmembrane domains (DI-DIV), each with six transmembrane segments (S1-S6), as shown in Fig. 1A. Segments S1-S4 constitute the voltage-sensing domains (VSDs), with the S4 helix containing multiple arginine residues crucial for sensing changes in the transmembrane voltage, while S5-S6 segments form the pore domain (PD), responsible for creating the central ion conduction pore(*27, 28*). Our analysis revealed that seven of the eight most common coding variants of *CACNA1A* reside in the cytoplasmic loop connecting the transmembrane domain I and II of hCa_V_2.1, whereas all *de novo* variants of *CACNA1A* are located in conserved transmembrane regions throughout the hCa_V_2.1 protein (Fig. 1A). Interestingly, nearly a third of *de novo* variants are located in the transmembrane S4-linker-S5 segment of transmembrane Domain III, underscoring the functional significance of this region.

**Fig. 1.**
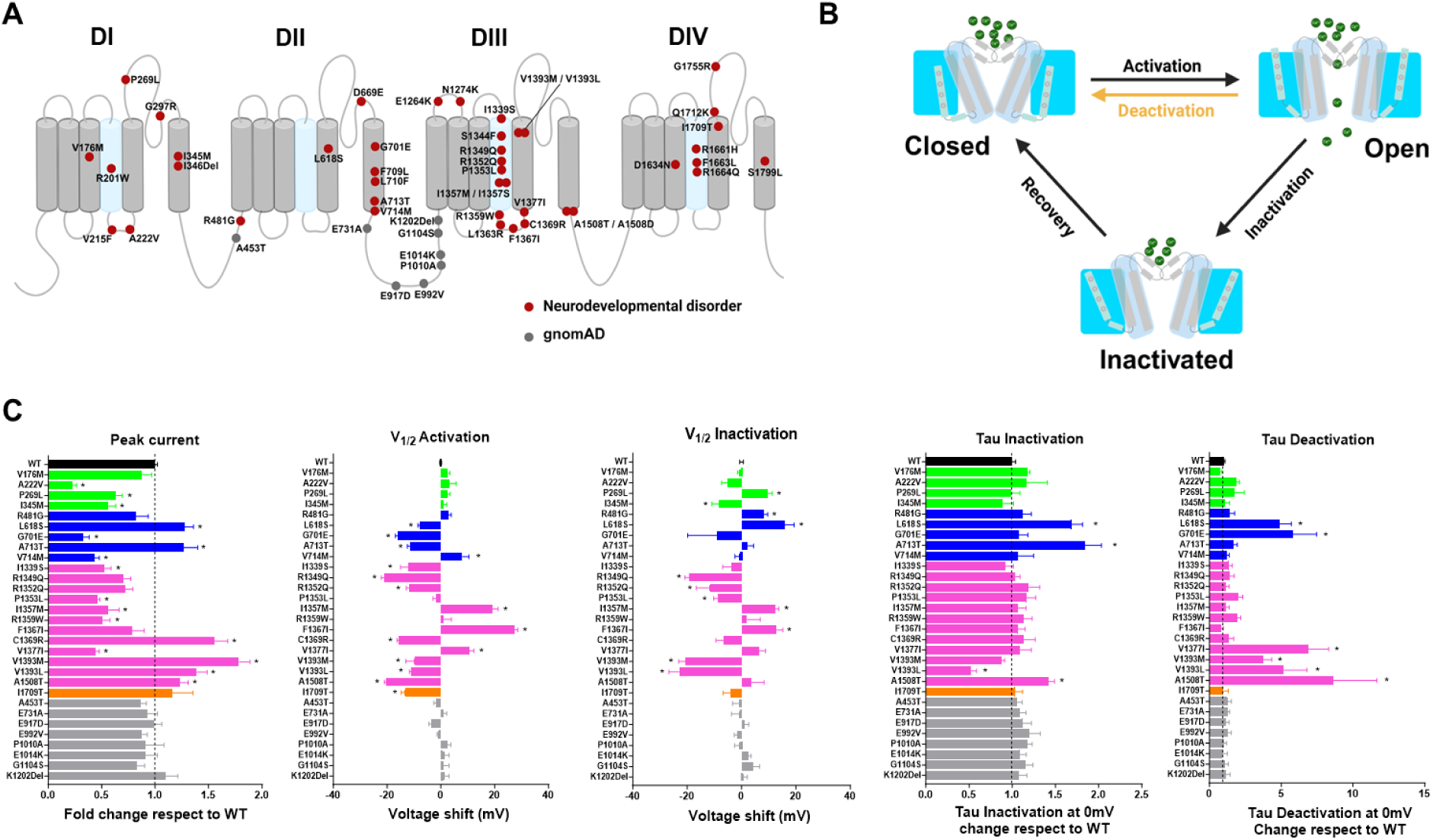
Functional characterization of *de novo* coding variants of *CACNA1A in patients with severe neurodevelopmental disorders*. **A:** Schematic representation of the membrane topology of the hCaV2.1 channel encoded by CACNA1A. DI-DIV corresponds to four transmembrane domains. The S4 segment is the voltage sensor and is shown in light blue in each domain. The location of the *de novo* missense from the cohort (red) and common variants from gnomAD (grey) are indicted on the membrane topology. **B:** The cartoons represent three major structural conformations of the hCa_V_2.1 channel: Closed, Open and Inactivated *states.* The transitions between the three conformations depend on the membrane voltage. In Closed state the voltage sensing domain (VSD) is in downward position and ions in the cavity are not permeated; Open state indicates that VSD is translocated to the upward position to allow ion conduction ions; Inactivated state shows that VSD is in a different upward configuration and the ions can not be permeated through the pore domain. **C:** Biophysical characterization of *de novo* and common missense variants of *CACNA1A*. Each bar represents one variant hCa_v_2.1 channel encoded by a missense change, and the graph is organized based on domain locations: DI variants in light green, DII in blue, DIII in magenta and DIV in orange while gnomAD variants in grey. “Peak current” shows the peak current densities for each variant hCa_v_2.1 channel, normalized to the wild-type channel. Changes in the peak current densities are shown as fold change values. Voltage-dependent channel activation, “V_1/2_ Activation”, shows the change in the V_1/2_ ACT (ΔV_1/2-ACT_) for each variant hCa_v_2.1 channel with respect to the wild-type channel (x=0). The presentation is similar for “V_1/2_ Inactivation”, where ΔV_1/2-INACT_ for each variant of hCa_v_2.1 channel is shown. x=0 in V_1/2_ Activation and V_1/2_ Inactivation bar graph corresponds to the wild-type (WT) hCa_v_2.1 channel behavior. For “Tau Inactivation’’, the time constant (τ) of inactivation decay for each hCa_v_2.1 variant channel is normalized to the wild-type channel (x=1). The similar presentation for “Tau Deactivation”, the time constant (τ) of deactivation decay for each hCa_v_2.1 variant channel is normalized to the wild-type channel (x=1). Asterisk denotes significant differences from the wild-type channel; see Table 1.

Ca_V_ channels exist in conformations that roughly correspond to *closed states, open states, and inactive states* (Fig. 1B)(*1, 29, 30*). The relative occupancy in open, closed, and inactive states defines how the channel responds to physiological stimuli in neurons(*31–33*). We established electrophysiological recording protocols (Materials and Methods) to probe the transition between the open (O), closed (C), and inactivated (I) states of hCa_V_2.1 channels by determining five key biophysical properties: voltage dependence of activation, voltage dependence of inactivation, peak current density, and time course of inactivation and deactivation (Suppl. Fig. 2). These biophysical properties produce a comprehensive functional landscape for each hCa_V_2.1 variant relative to the WT channel. To measure these properties, we transiently transfected WT or variant hCa_V_2.1 cDNA into the Ca_V_β_3_-expressing cells, recorded variant and WT channel currents using whole cell voltage clamp as we previously described(*20*), and analyzed the function of variant channels compared to WT.

### *De novo* variants of *CACNA1A* alter many aspects of the channel properties Peak current density

Unexpectedly, we were not able to detect significant currents in 20 (of 42, ∼47.6%) hCa_V_2.1 variant channels encoded (Δfold change > −0.95, Suppl. Fig. 3). The majority (30 of 42, 71%) of hCa_V_2.1 variant channels exhibited a significant reduction in the whole-cell current density compared to WT channels. In contrast, hCa_V_2.1 channels encoded by each of the eight common variants displayed a similar amplitude of current density compared to WT channels (Fig. 1A “Peak current*”*, Peak current density in Table 1). Interestingly, six *de novo* missense changes resulted in significantly increased current density, most notable was variant V1393M (Δfold = 1.78 ± 0.1, *p* <0.0001). Only six *de novo* coding variants (V176M, R481G, R1349Q, R1352Q, F1367I, and I1709T) among the 42 did not significantly alter Ca_V_2.1 current density. These results are somewhat surprising, as the loss of function of *CACNA1A* was not previously known to be a strong genetic risk for neurodevelopmental disorder in cohort studies that focused on protein-truncating variants. Our results showed that the majority of the *de novo* variants of *CACNA1A* identified in patients with neurodevelopmental disorder greatly reduced the current density in the heterologous expression system, indicating that this property may underlie significant risk in neurodevelopmental disorders.

### Voltage-dependent activation

Of the 22 *de novo* variants that produced measurable Ca_V_2.1 currents, nearly three-quarters (15) produced hCa_V_2.1 channels opening at membrane voltages significantly different from WT hCa_V_2.1 channels (Fig. 1C, “V1/2 Activation”, V1/2 Act in Table 1). Among these 15 variants, 11 shifted the voltage dependence of activation to more hyperpolarized potentials (left shift in V_1/2-ACT_), while 4 shifted to more depolarized potentials (right shift in V_1/2-ACT_). The left shift of V_1/2-ACT_ indicates increased calcium influx during action potentials, whereas the right shift of V_1/2-ACT_ reduces calcium influx. The voltage dependence of activation of hCa_V_2.1 channels coded from any of the eight common variants was indistinguishable from that of WT hCa_V_2.1 channels (Fig. 1C, grey bars, Table 1). Only one *de novo* variant (A1508T, k slope = 4.1 ± 0.9 compared to 7.9 ± 1.2 in WT) exhibited a difference in the steepness of the channel activation voltage dependence (Table 2). Notably, hCa_V_2.1 channels coded by A1508T opened at the most negative membrane voltages relative to those of the WT hCa_V_2.1 channels (ΔV_1/2-ACT_ = - 20.3 ± 0.7 mV, p<0.001), while F1367I produced channels opened at the most positive voltage (ΔV_1/2-ACT_ = 27.2 ± 1.3 mV).

**Table II.**
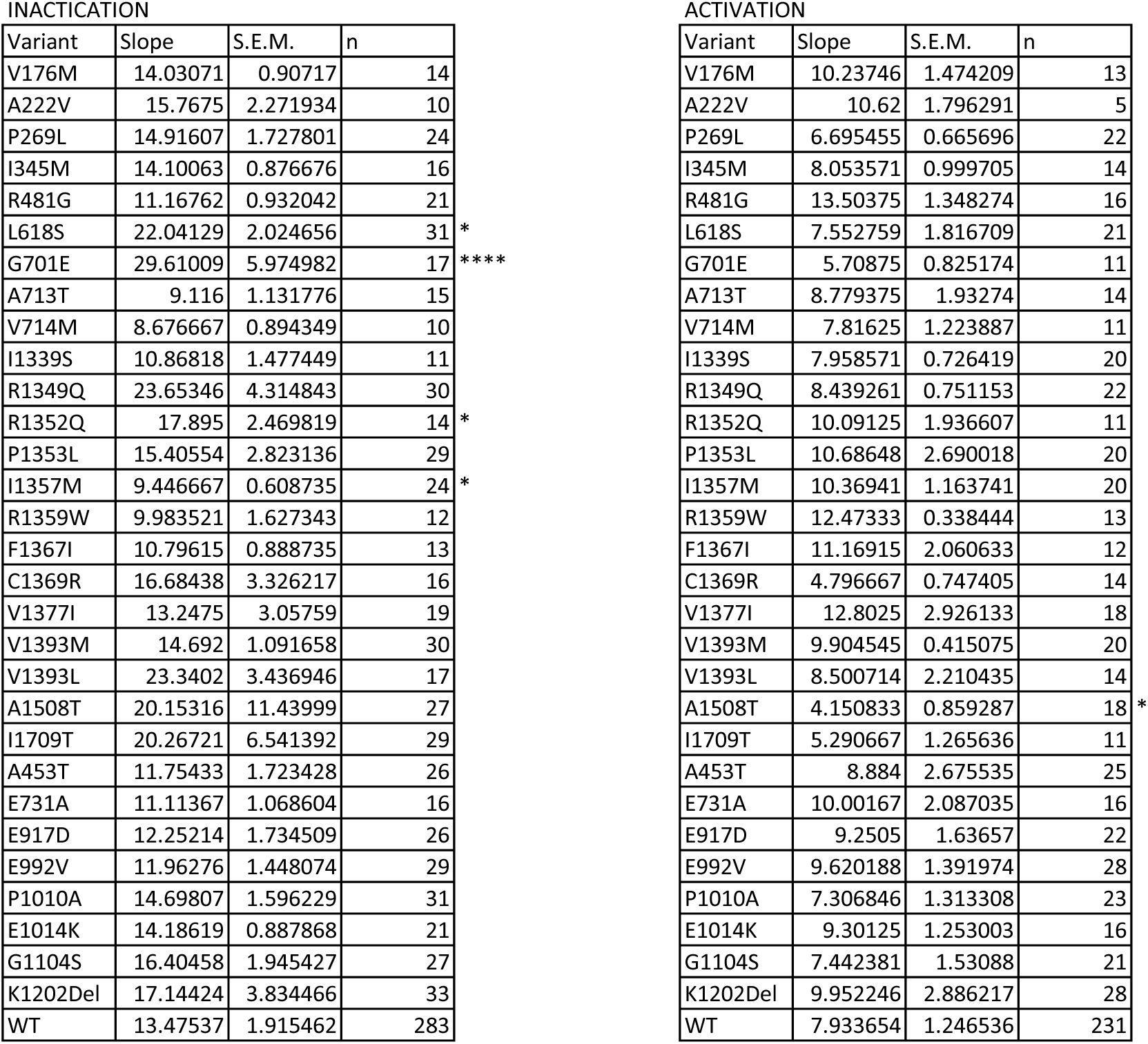
Slope for Boltzmann fit for voltage dependent inactivation and activation of the variants from Neurodevelopmental disorders and gnomAD cohorts. Twenty variants without currents were excluded from our analyses.

In summary, 15 of 22 (68%) *de novo* variants significantly altered how channel responds to membrane depolarization, as defined by the V_1/2-ACT_ values relative to the WT, while none of the common variants did, suggesting that the voltage-dependent activation of the hCa_V_2.1 channel may be specific to disease-associated phenotypes.

### Voltage-dependent inactivation

Out of 22 *de novo* variants measured, 11 (50%) altered the voltage dependence of inactivation (V_1/2-INACT_) compared to the WT channel. Six shifted V_1/2-INACT_ towards hyperpolarized potentials, while five shifted it towards depolarized potentials (Fig. 1C, “V1/2 Inactivation”, V_1/2-INACT_ in Table 1). Notably, V1393L exhibited the most negative shift (ΔV_1/2-INACT_ = –23.0 ± 3.8 mV, *p* < 0.0001) compared to WT hCa_V_2.1, whereas L618S channels were inactivated at the most depolarized potentials (ΔV_1/2-INACT_ = 16.0 ± 3.3 mV, *p* < 0.0001). These analyses revealed that 50% of the *de novo* variants and none of the common variants produced channels with inactivation voltages differing from WT hCa_V_2.1, suggesting that the voltage-dependent inactivation is a significant biophysical property associated with human coding variants associated identified in neurodevelopmental disorders. Interestingly, we found that V_1/2-Act_ and V_1/2-Inact_ are significantly correlated (p=0.0006, r2=0.34), indicating that variants induced changes in sensing voltage similarly impact activation and inactivation.

### Kinetics of deactivation and inactivation

We found that 6 of 22 (27.3%) hCa_V_2.1 channels produced by *de novo* variants (L618S, G701E, V1377I, V1393M, V1393L, and A1508T) significantly slowed the rate of deactivation relative to WT channels, while no common variants altered this property in hCa_V_2.1 channels (Fig. 1C, “Tau Deactivation”, τ deact in Table 1). The variant with the slowest rate of deactivation in our dataset was A1508T (Δ τ _DEACT_ = 8.6 ± 3.0, *p* <0.0001) compared to the WT baseline (Δ τ _DEACT_ = 1 ± 0.8).

Similarly, we found 4 of 22 (18.2%) hCa_V_2.1 channels produced by *de novo* variants significantly altered the rate of inactivation relative to WT channels (L618S, A713T, V1393L, and A1508T). The channel with the slowest rate of inactivation was A713T (Δτ_INACT_ = 1.8 ± 0.2, *p* <0.0001, compared to WT Δτ_INACT_ = 1 ± 0.04). Interestingly, the V1393L variant showed a significantly faster inactivation rate compared to WT (Δτ_INACT_ = 0.52± 0.05, p<0.005). This change did not dependent on the change in the voltage dependence of activation in V1393L channels, as we compared the τ_INACT_ over a range of voltages. All changes in kinetics induced by coding variants were accompanied by change in voltage dependence of activation and/or inactivation, suggesting that these structural positions are highly sensitive to multiple channel biophysical properties.

In summary, nearly half (20 out of 42) of *de novo* coding variants in this allelic series of *CACNA1A* produced hCa_V_2.1 channels without detectable currents. Of the 22 *de novo* variants (of 42) that produced measurable hCa_V_2.1 channels, ten encoded channels with a significant reduction in current density compared to reference WT channels, 15 altered the voltage dependence of hCa_V_2.1 channel activation, and 11 resulted in channels with an altered voltage dependence of inactivation. Additionally, six *de novo* variants slowed the time course of deactivation, and four altered the inactivation kinetics. None of these changes were observed in common variations. All but one of the *de novo* variants produce hCa_V_2.1 channels that were significantly different from WT in at least one aspect of examined biophysical properties. These results demonstrate that *de novo* coding variants of *CACNA1A* drastically impair channel properties in various functional domains, altering calcium influx, and strongly implicate *CACNA1A* in neurodevelopmental disorders where these *de novo* variants were identified.

### *De novo* variants that alter channel peak current density and voltage dependence of activation are located in distinct structural regions

To visualize and compare the functional impact of coding variants across multiple biophysical properties, we calculated the functional differences among hCa_V_2.1 variants relative to WT channels with dimensionless z-scores. For this analysis, we included the 22 variants that produced measurable Ca^2+^ currents (Table 1), with z-scores calculated for each of the five biophysical properties of each variant channel (Methods, Suppl. Table 1). Using z-scores, we visualized the overall functional impact of each *de novo* missense variant by radar plot, where each corner represents a distinct biophysical parameter (Fig. 2, inset). As shown in Fig. 2A, z-scores calculated across the five electrophysiological parameters were plotted on radar maps based on the topological location (Domain I to Domain IV), while gnomAD variants, all located in intracellular domains, were shown in grey pentagons. A statistical comparison of disease-associated variants in the four domains of Ca_V_2.1 to gnomAD variants for each functional property is shown in Fig. 2B. Variants from transmembrane domain III (D3) as a group were significantly different in peak current density (p = 0.0048), V½ activation (p = 0.0022), and V½ inactivation (p = 0.0043) compared to gnomAD variants. Similarly, *de novo* variants in Domain II (D2) as a group also show significant differences in peak current density (p = 0.041) and V½ activation (p = 0.036), compared to gnomAD variants. Interestingly, only one (of 8 *de novo*) variant in Domain IV (D4) appeared in this plot because we could not reliably detect the current density for the other seven. Together, these results demonstrated that peak current density and V½ activation and inactivation are most susceptible to *de novo* coding changes in hCa_V_2.1 channels. In addition, different VSD domains mediate distinct changes - *de novo* variants in Domain IV mainly altered peak current density while those in Domain II/III altered multiple biophysical changes of the channel, consistent with the notion that different channel structural elements play distinct roles in channel function.

**Fig. 2.**
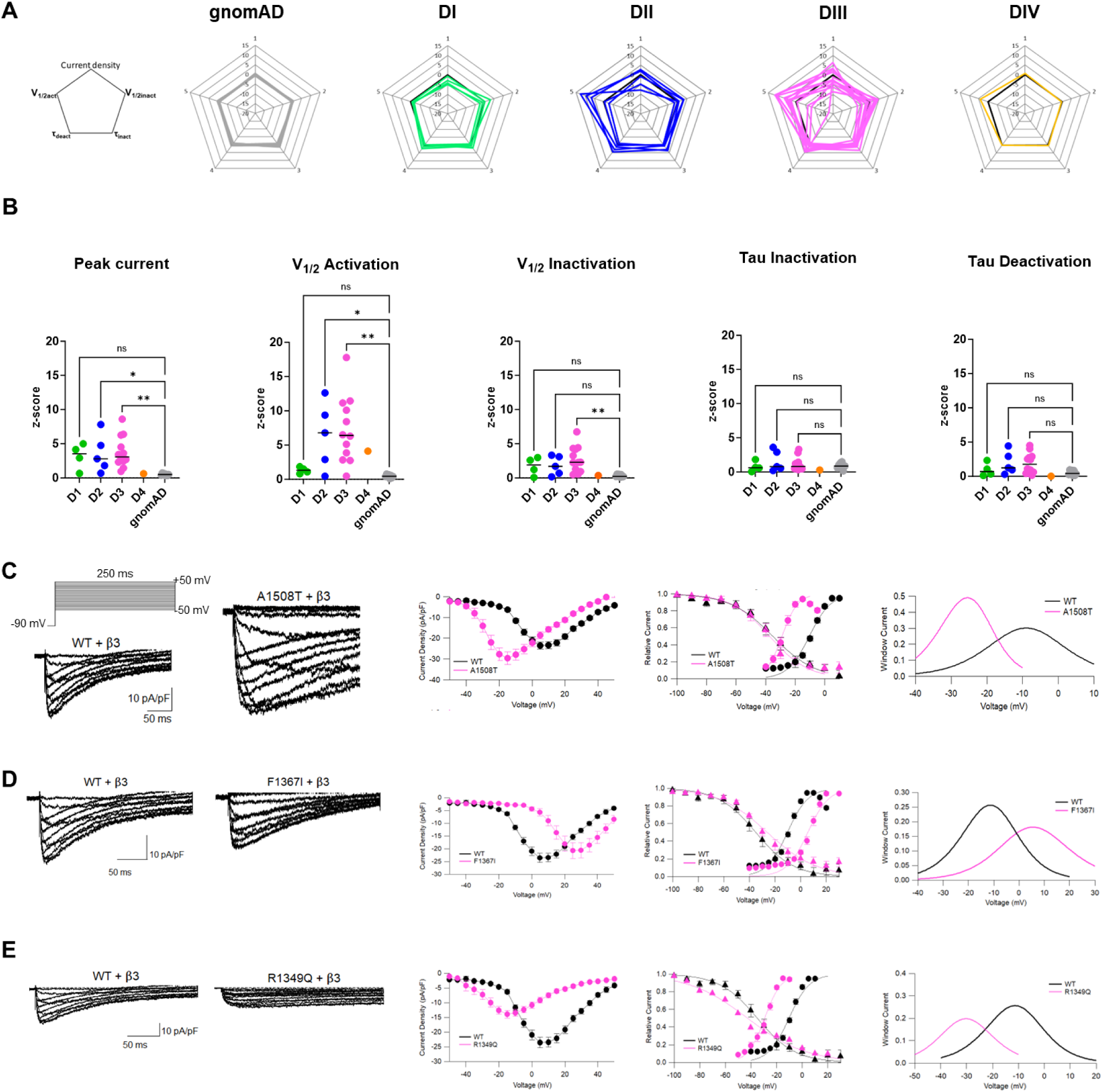
Functional comparison among *de novo* missense variants and variants from gnomAD in *CACNA1A* using z-scores and sample traces of hCav2.1 variants. **A,** The five biophysical properties for each variant hCa_v_2.1 channel with detectable currents are visualized using dimensionless z-scores in a pentagon. The data shown here corresponds to 30 of the 50 variants presented in Figure 1A and does not include those without measurable currents. Grouped pentagon from gnomAD variants (grey) and *de novo* variants in four structural domains (DI: light green, DII: blue, DIII: magenta and DIV: orange) are shown. Each pentagon represents a variant, where each corner corresponds to the z-core of one biophysical parameter (left inset). The solid black line in each rhombus plot indicates the wild-type behavior. The z-score for wild-type is 0. **B,** Domain-level comparison of absolute z-scores for each of the five properties of the hCa_v_2.1 channels. Absolute z-scores for *de novo* variants in each domain are compared to those of gnomAD. C-E, *Left*: Raw traces of hCa_v_2.1 currents from cells expressing WT, A1508T (top, **C**), F1367I (middle, **D**) and R1349Q (bottom, **E**) in the presence of β_3_ subunit. Currents were evoked by 250 ms long test potentials from a holding potential of −90 mV. *Right*: Average current-voltage plots, voltage-dependence of activation and inactivation and window currents for WT, A1508T (top, **C**), F1367I (middle, **D**) and R1349Q (bottom, **E**). Solid line in voltage-dependent activation and inactivation was fit with Boltzmann equation. The window current was estimated by multiplication of activation and inactivation curve from Boltzmann function (see method).

Fig. 2C-2E showed the sample traces of whole cell clamp recording of the variant’s channels, indicating increased channel current density (Fig. 2C) and rightward (Fig. 2D) and leftward (Fig. 2E) shifts in voltage-dependence of channel activation. With Ba^2+^ as the charge carrier in whole-cell recordings, cells expressing WT hCav2.1 channels showed, and average barium currents peaked at 4.8 ± 0.6 mV with a current density of 22 ± 10.7 pA/pF. The current densities in cells expressing A1508T, but not F1367I, were larger relative to those in cells expressing WT hCa_v_2.1, while the R1349Q variant exhibited reduced current density. In addition to the current density, cells expressing A1508T and R1349Q displayed large hyperpolarized shifts (ΔV_1/2-ACT_ = −20.3 ± 0.7 mV and −20.9 ± 1.2 mV, respectively), while F1367I exhibited a large 27.2 ± 1.3 mV depolarized shift in the voltage-dependence of channel activation (Fig. 2C, D and E pink) in comparison to WT. Moreover, the calculated window currents for A1508T and R1349Q variants, but not F1367I, showed a hyperpolarized shift in voltage range. These data illustrate how missense variants in hCa_V_2.1 can induce drastic functional effects. To gain additional structural insight on the impact of coding variants, we mapped the variants to the 3D channel structure, focusing on those variants that either altered the peak current density or the voltage dependence of activation.

### 3D structural modeling revealed structural hotspots for missense changes in current density and voltage dependence of activation

Based on the modeled structure of hCa_V_2.1 using AlphaFold 2 (Fig. 3A, method), we were able to map all 27 *de novo* coding variants that showed either no current or reduced currents (see method, Fig. 3A, R201W, V215F, A222V, G297R, I346del, D669E, G701E, F709L, L710F, E1264K, N1274K, I1339S, S1344F, P1353L, I1357S, R1359W, L1363R, F1367I, V1377I, A1508D, D1634N, R1661H, F1663L, R1664Q, Q1712K, G1755R and S1799L) in the channel 3D structure. To better visualize the structural elements, we collapsed the four VSD domains and aligned them to domain I. *De novo* variants that induced significant current density reduction roughly mapped to three regions (Fig. 3B). Several of the variants exhibiting low current density (12 of 27; 44%) are located in Region I, corresponding to S5-S6 segments in the pore domain (PD). Of the 12 variants, four residues (G297R, D669E, Q1712K and G1755R) are located on the extracellular loop between S5-S6 segments, with 2 variants are close to the selectivity filter, which could be involved in either obstructing the channel opening or repelling Ca^2+^ ions. The remaining 8 variants (A222V, I346Del, G701E, F709L, L710F, V1377I, A1508D and S1799L) are located on the transmembrane S5 and S6 segments (Fig. 3C). Region II refers to the S4 segments and S4-S5 linkers, where 10 variants are mapped (R201W, V215F, P1353L, I1357S, R1359W, L1363R, F1367I, R1661H, F1663L and R1664Q). Among variants mapped to this region, four directly involved voltage sensing arginine residues (R201W, R1359W, R1661H, and R1664Q). Region III encompasses the extracellular S1-S2 and S3-S4 linkers in the VSD domains (E1264K, N1274K, I1339S and S1344F). These analyses highlighted the PD domain, S4 segments, and the S4-S5 linkers as hot spot regions where coding variants affect channel function.

**Fig. 3.**
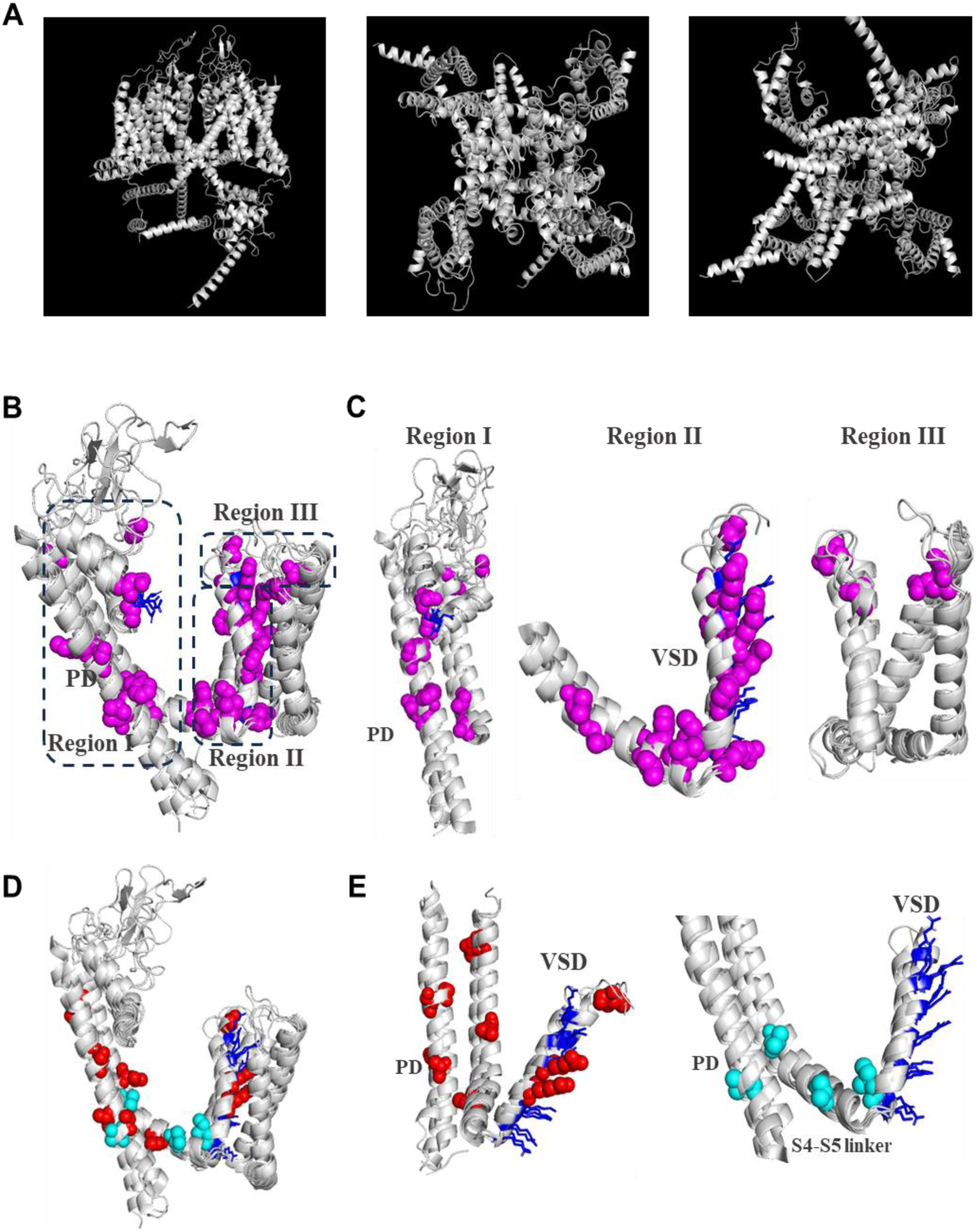
3D modeling revealed structural hotspots for missense changes that altered channel current density and voltage-dependent activation. **A,** Side view (left), top view (middle) and bottom view (right) of the structural model of hCa_v_2.1 obtained from Alphafold2 database with optimizations. **B,** The four transmembrane domains (DI-DIV) are structurally aligned to DI, and the residues that altered current density are indicated in magenta. Aligned transmembrane domains display three hotspot regions. **C,** Three regions (from **3B**) are shown separately with variants that altered current density. The positively charged arginine residues in S4 segment and selectivity filter residues for Ca^2+^ selectivity are highlighted in blue. **D,** Fifteen variants that changed voltage-dependence of activation are mapped onto the transmembrane domain alignment of hCa_v_2.1 that highlighted S4 segment and S5-S6 pore region. **E,** Variants shifted voltage-dependent activation to the left (in red) and right (in cyan) (from **3D**) are shown separately in S4 segment and S5-S6 pore region.

We next mapped the structural location of the 15 variants that altered hCa_V_2.1 voltage-dependent activation (V_1/2-ACT_) (Fig. 3D). Out of the 15 variants, 11 exhibited a hyperpolarized shift (Fig. 3D, red residue, L618S, G701E, A713T, I1339S, R1349Q, R1352Q, C1369R, V1393M, V1393L, A1508T and I1709T), while the remaining 4 variants shifted the voltage dependence of activation to more depolarized voltages (V714M, I1357M, F1367I and V1377I in cyan). Located in the S4 segment, R-to-Q mutations (R1349Q and R1352Q) did not affect current density, but shifted voltage-dependent activation to more hyperpolarized potentials (ΔV_1/2-ACT_= −20.9 +1.2 mV for R1349Q and ΔV_1/2-ACT_= −11.9 +0.9 mV for R1352Q), consistent with the notion that these arginine residue plays a key role in voltage-sensing (Fig. 3E, red residues). Seven of the 11 mutants that caused hyperpolarized shifts in V_1/2-ACT_ are located within transmembrane S5 or S6 segments. For example, V1393M, V1393L and I1709T are located on the top of the S5 segments of DIII and DIV, respectively. All three variants shifted the activation curve to more negative potentials (ΔV_1/2-ACT_= −9.8 +3.2 mV for V1393M, ΔV_1/2-ACT_= −11.0 +0.6 mV for V1393L and ΔV_1/2-ACT_= −13.4 +1.4 mV for I1709T), highlighting the role of this region in transducing voltage changes to pore opening. Interestingly, although two A-to-T variants located on the bottom of S6 segment exhibited a strong hyperpolarized shift in V_1/2-ACT_ (ΔV_1/2-ACT_= −11.5 +0.9 mV for A713T and ΔV_1/2-ACT_= −20.3 +0.7 mV for A1508T), a variant (V714M) affecting an adjacent residue shifted the activation curve to more depolarized potentials (ΔV_1/2-ACT_= 8.0 +2.5 mV) (Fig. 3D and Table 1), revealing contributions of this microregion in mediating the voltage dependence of activation. Three out of four variants (I1357M, F1367I and V1377I) that shifted the activation curve to more depolarized potentials are all located on the S4-S5 linker (Fig. 3E, cyan residues), and the adjacent region of the S5 segment, respectively. While V1377I induced a +10 mV shift (ΔV_1/2-ACT_= 10.5 +1.7 mV), the other two variants caused +20 and +27 mV shifts (ΔV_1/2-ACT_= 19.1 +2.1 mV for I1357M and ΔV_1/2-ACT_= 27.2 +1.3 mV for F1367I), respectively, consistent with a role of the S4-S5 linker in transducing voltage sensing to pore opening. Similarly, we mapped the *de novo* variants that shifted the voltage-dependence of inactivation, many of which exhibited concurrent changes in the current density and voltage-dependence of activation (Suppl. Fig. 4).

In summary, the *de novo* coding variants in *CACNA1A* affect channel function by mainly altering the overall current density and the channel responses to membrane potentials. We highlighted specific microregions that are hotspots for these functional effects. While these coding variants are spread across many domains, but variants in S4, S4-S5 linker, and S5-S6 pore regions are impactful.

### *NEURON* simulation reveals the impact of the biophysical properties of the hCa_V_2.1 channel on Purkinje neuron excitability

Ca_V_2.1 channels play a crucial role in Purkinje neuron excitability, essential for cerebellum function(*34, 35*) which may underlie *CACNA1A* disease pathophysiology(*5, 9*). To understand the physiological impact of biophysical changes of hCa_V_2.1 channels, we simulated neuronal excitability using a mathematical model of a multi-compartment Purkinje neuron deposited in *NEURON*(*23*) (ModelDB, 229585), with a complex arborization of 1600 dendrites (Fig. 4B) (Suppl. Fig. 5)(*18, 36*), and ionic conductance for Na^+^, K^+^ and Ca^2+^. Using this model, we discovered that altered current density (CD) and the voltage dependence of activation of hCa_V_2.1 channels (V_1/2-ACT_) affected both the number of action potentials produced and the latency to the first action potential in spontaneous firing (Fig. 4A, I-IV). The firing activity generated by the model in the presence of WT hCa_V_2.1 channels is characterized by a monotonic spontaneous firing of ∼0.02 Hz (54 spikes/s) and a latency of ∼290 ms (Fig. 4A I). Altering the biophysical properties of the hCa_V_2.1 conductance leads to longer latency and lower firing rate (II), or irregular bursting-like complex firing patterns followed by a long period of inactivity (III), including a regular busting phenotype (IV). Based on our simulations, changes in hCa_V_2.1 current density drastically modified the latency to the first action potential, ranging between ∼900 ms to ∼200 ms with 35% to 200% of the WT value, respectively (Fig. 4C, Top). By contrast, changes in current density did not affect the firing rate significantly until there is only ∼35% of total current density left and then the neuron stopped firing (Fig. 4C, Bottom). In comparison, a shift in the V_1/2-ACT_ to more negative voltages greatly reduced the latency to the first action potential, changing from 288 ms in WT condition to 56 ms when V_1/2-ACT_ is shifted −20 mV. The rightward shifts can increase the latency to the first action potential to ∼900 ms, and when the V_1/2-ACT_ is shifted more than +4 mV with respect to WT, the system is unable to fire (Fig. 4D). Interestingly, changes in the firing rate induced by changes in hCa_V_2.1 V_1/2-ACT_ have three flavors. When ΔV_1/2-ACT_ is between −4 and +4 mV from WT, the spontaneous firing remains WT-like tonic firing pattern (100 spikes/sec at −4 mV, to 45 spikes/sec at +4 mV, respectively). When ΔV_1/2-ACT_ left shifted more than 5 mV, the firing pattern becomes complex bursting (Fig. 4A. III and 4A. IV) (Suppl. Fig. 5). When ΔV_1/2-ACT_ is more positive than 4 mV, the system is not able to produce any action potentials (Fig. 4D). The overall impact of current density and V_1/2-ACT_ on the spontaneous Purkinje output is summarized in Fig. 4C-E.

**Fig. 4.**
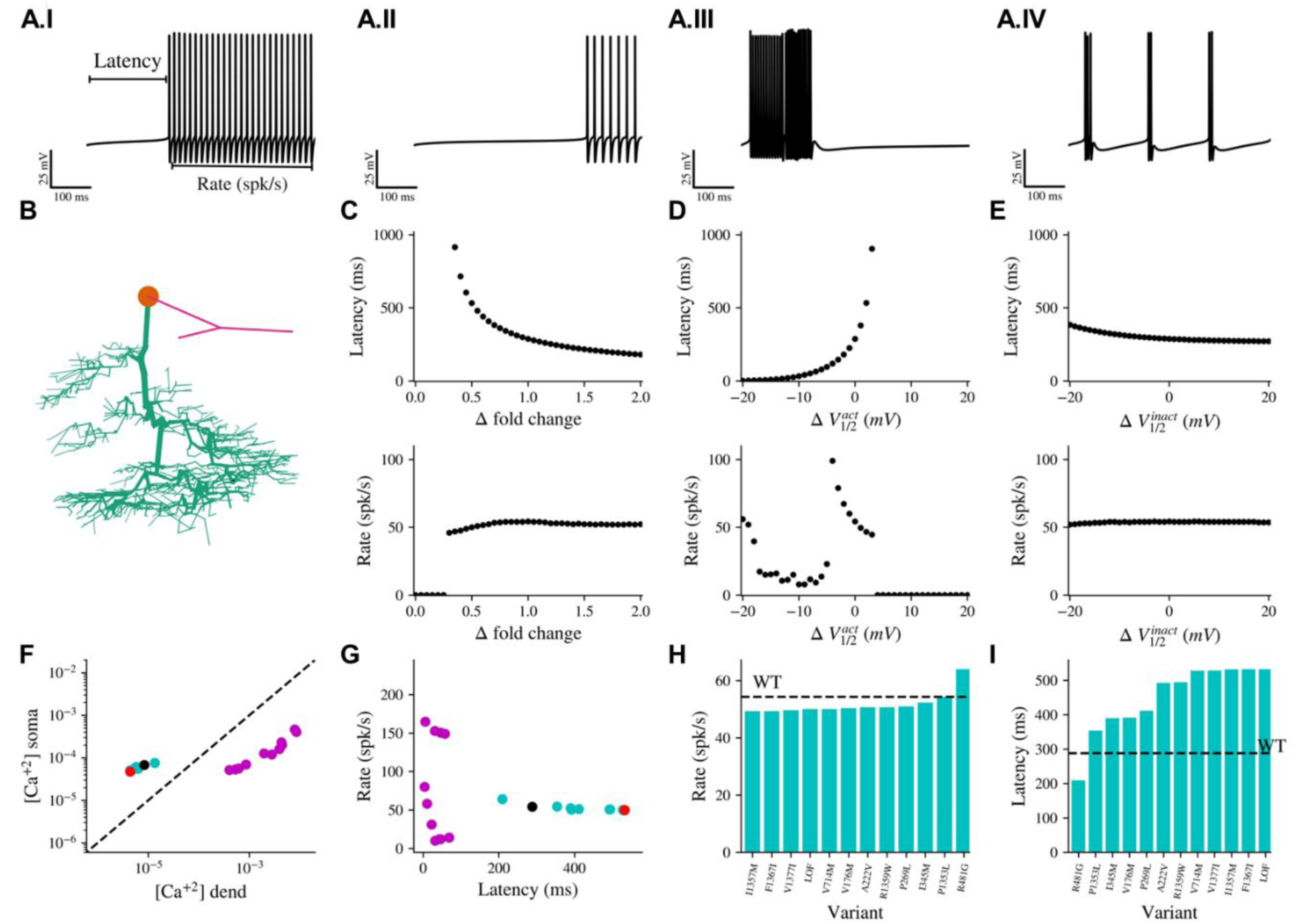
Impact of the biophysical properties of the hCa_v_2.1 variants on NEURON Purkinje cell model reveal drastic changes on firing properties. **A.** Sample of simulated axonal voltage traces of spontaneous firing from a Purkinje cell model using *NEURON* showing different types of activity when hCa_v_2.1 properties were modified. (I) Default conditions WT, II-IV changes in the firing associated with changes in the biophysical properties of the hCa_v_2.1 conductance. **B.** Spatial organization of the NEURON model (DB 229585, see methods). Dendritic arborization is shown in green corresponding to 1600 of dendrites, soma is represented in orange, and the axonal compartments in magenta. **C.** Impact of hCa_v_2.1 CD on the spontaneous activity firing properties. **D.** Impact of changes in hCa_v_2.1 voltage activation midpoint on the spontaneous activity firing properties. **E**. Impact of changes in hCa_v_2.1 voltage inactivation midpoint on the spontaneous activity firing properties. **F.** Relation between Ca^2+^ measurement at the dendrites and soma for all the variants simulated, WT conditions are represented in black, and heterozygous LoF expression is represented in red, dash line showing the diagonal. **G.** Correlation between “Latency” and “Firing rate” from the simulated spontaneous firing activity for all the variants characterized in this study, WT conditions are represented in black, and heterozygous LoF expression is represented in red. **H**. Spiking rate (#spikes/s) of the spontaneous activity simulated for all the variants that presented a tonic firing, black dotted line represents WT conditions (54 spikes/s). **I.** Latency time of the spontaneous activity simulated for all the variants that presented a tonic firing, black dotted line represents WT conditions (290 ms). All the simulations were run for 5 seconds, the calcium concentration in F corresponds to the average of the final 3 seconds (see methods for more details).

Interestingly, the impact of the voltage dependence of inactivation (V_1/2-INA_) of hCa_V_2.1 on Purkinje excitability (Fig. 4E) is noticeable yet modest in the latency compared to that of current density or V_1/2-ACT_, and it did not have any major effect on the firing rate, suggesting that voltage-dependent inactivation of the channel does not play a major role in the simulated neuronal firing properties. Similarly, the impact of deactivation and inactivation kinetics is modest in this simulation (not shown).

Many *de novo* variants induced multiple biophysical property changes. For example, G701E and I1339S reduced current density by 70%, while it also shifted the activation threshold of the channel ∼20mV in hyperpolarized direction, which is consistent with more barium influx upon depolarizations. We thus analyzed the physiological impact of all *de novo* variants considering all the biophysical properties analyzed and in the presence of 50% WT channels to model heterozygosity. We found that we can categorize the variants into tonic firing and complex firing based on their excitability patterns. In addition, tonic firing (turquoise) and complex firing (fuchsia) variants can be easily distinguished by plotting the Ca^2+^ concentrations in soma as a function of the Ca^2+^ concentration in the dendrites (Fig. 4F). Variants that showed tonic firing clustered to the left of the diagonal (higher Ca^2+^ concentration in the soma), including the WT and heterozygous LoF (black and red dot respectively), while variants with a complex bursting firing pattern clustered together to the right of the diagonal (higher Ca^2+^ concentration in the dendrites). We then investigated the relationship between latency and action potential firing rates, without exception all the variants that shifted V_1/2-ACT_ to more negative potentials rendered shorter latency times (fuchsia), and complex firing patterns (Fig. 4A.III-IV/4D-Bottom, Suppl. Fig. 5). Despite a greater influx of dendritic Ca^2+^, these variants did not consistently show high firing frequencies. Instead, most of them presented lower or similar firing frequencies compared to WT (black dot), owing to complex firing patterns with long inter-burst periods. For those variants that produced tonic firing (Fig. 4AI-II), during the steady state at the end of the five-second simulations, differences in the firing rate were modest (Fig. 4G). However, some greatly affected the latency to the first action potentials, where the most extreme case was the heterozygous LoF (50% of the total WT Cav2.1 conductance) represented in red (Fig. 4G). Interestingly, when we look only at the variants that produced tonic firing, it becomes clear that some of these biophysical changes resulted in modest changes in the spike firing rate, with the major impact on the latency (Fig. 4H-4I). After simulation, we quantified physiological impact of *de novo* coding variants of *CACNA1A* on Purkinje output that include 1) pattern - tonic or complex firing; 2) latency - the time it takes to the first action potential in this simulation, 3) Mean steady state concentration of Ca in soma; 4) Mean steady state concentration of Ca in distal dendrites (Suppl. Table 2). We subsequently used these four neuronal outputs to correlate with clinical phenotypes (see Fig. 6).

### Coding variants of *CACNA1A* increase disease burden in developmental disorder and epilepsy using AlphaMissense Prediction

After functionally annotating 42 *de novo* coding variants of *CACNA1A*, we sought to extend our study of *CACNA1A* missense variants to other disease cohorts by evaluating computational prediction algorithm to understand the disease burden. Alpha Missense (AM) was developed to predict the pathogenicity of missense coding changes based on structure and sequence information together with the frequency in the population using a deep learning model(*37*). AM predicted the vast majority (39/41=95.1%) of analyzed *de novo* variants to be pathogenic, while one of the eight (12.5%) gnomAD common variants was predicted to be pathogenic (Fig. 5A, left two columns). This is largely consistent with our functional studies with > 85% accuracy. When extending the analysis to the full set of 659 missense variants of *CACNA1A* in gnomAD, we reinforced the substantially higher pathogenic coding mutation rate of *CACNA1A* in the neurodevelopmental disorder cohort than in the general population (Fig. 5A, right). Moreover, utilizing a recent exome-sequencing study of epilepsy, which comprises ∼21,000 epilepsy cases with clinical phenotypes and ∼33K well-matched controls(*24*), we found elevated pathogenic mutation rates across all epilepsy subtypes compared to the controls, with the most significant and strongest burden observed in the most severe subtype - developmental and epileptic encephalopathy (DEE), which is a group of epilepsies characterized both by seizures and severe developmental delay (Fig. 5B). Collectively, these results suggest a significant role of *CACNA1A* missense variants in epilepsy in general and increasing risk for DEE in particular. Similar analyses of the protein truncating variants did not show significant disease burden, underscoring the role of the missense change of *CACNA1A* in neurodevelopmental disorder susceptibility. Notably, AlphaMissense prediction does not correlate with any of the five physiological measurements of mutant Ca_V_2.1 channels (Fig. 5C), suggesting that AlphaMissense is unable to predict the directionality or impact on channel functional induced by missense alterations. This pattern highlights the value of sensitive functional assays in providing valuable information on the molecular impact of missense variants, facilitating the identification, characterization and interpretation of disease-associated mutations that has not been achievable by in silico predictions.

**Fig. 5.**
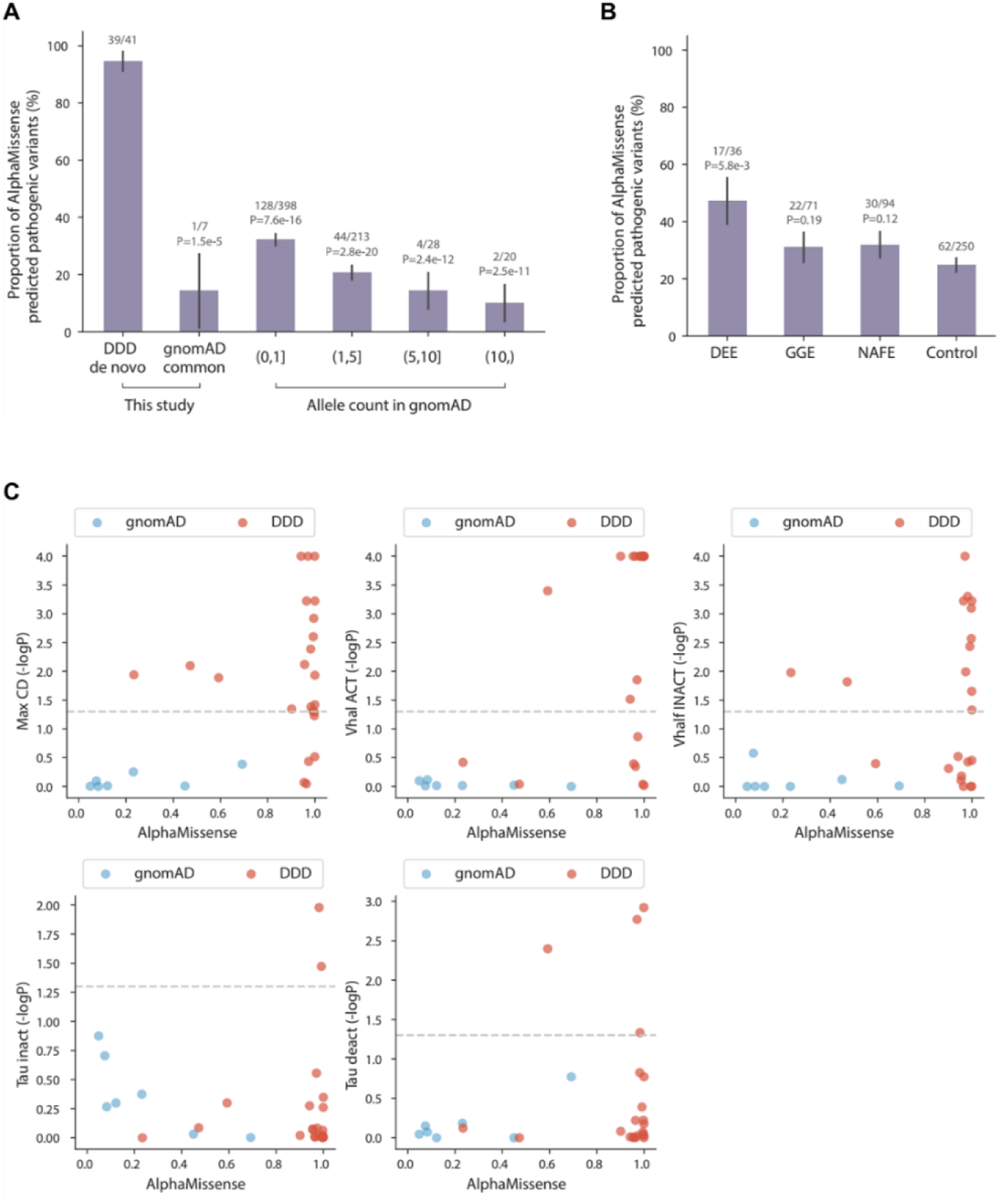
Predicted pathogenic missense variants of *CACNA1A* increase disease burden in developmental disorder and epilepsy. **A,B**. Proportions of missense variants predicted to be pathogenic by AlphaMissense, categorized by disease cohort - *de novo* variants found in individuals with neurodevelopmental disorder (DDD), population variants reported in Genome Aggregation Database (gnomAD) with no known neurodevelopmental disorders (**A)**; rare variants found in individuals with epilepsy across three subtypes: developmental and epileptic encephalopathy (DEE), genetic generalized epilepsy (GGE), and non-acquired focal epilepsy (NAFE), as well as rare variants in controls sequenced in the same epilepsy WES study (**B**). Error bars indicate standard errors of the proportions. *P*-values are calculated using a Fisher’s exact test. **C**, Comparisons between AlphaMissense predictions and biophysical properties measured in this study of missense variants’ effects on 50 variant channels in more than 1000 cells. Missense variants found in individuals with DDD and in gnomAD are colored by red and blue, respectively. Dashed horizontal lines indicate significant (*P*-value<0.05) biophysical property changes induced by specific missense variants.

### The relationship between channel function and clinical phenotypes

Coding variants of Ca_V_2.1 have been reported in patients with a spectrum of disorders, including epilepsy, developmental delay, ataxia, and hemiplegic migraine. However, it is unclear whether certain molecular properties of Ca_V_2.1 channel are associated with specific clinical outcomes. We utilized two clinical datasets to address this question: one is based on the clinical information derived from the ∼31,000-trio cohort where most of the *de novo* variants of *CACNA1A* in this study originated, and the other one is based on a review of more than 300 *CACNA1A* variants reported in the literature or partnering clinical sites (see methods). From the information derived from the cohort, we mapped the 18 most frequent clinical phenotypes for 36 patients who each harbor one *de novo* missense change in *CACNA1A*. In comparison, we obtained 12 clinical phenotypes from 81 patients in the review study.

We utilized three measured biophysical properties (current density, V_1/2-ACT_, and V_1/2-INACT_) and four physiological readouts derived from simulated Purkinje firing for each *de novo* variant in heterozygosity to perform the analyses. The significance (-logP) for logistic regression analyses between seven metrics in molecular and neuronal function and the 18 clinical phenotypes derived from the cohort study (dataset one) is shown in Fig. 6A. We found that six (of the 18) phenotypes (hypotonia, seizure/EEG abnormality, abnormality of face, brain atrophy and gross motor delay) are significantly associated with at least one biophysical parameter. Notably, current density is significantly associated with hypotonia, brain atrophy, seizure/EEG abnormality, and abnormality of face, while V_1/2-ACT_ is significantly associated with hypotonia. For the physiological output, we found variants that produce tonic firing is significantly associated with brain atrophy, hypotonia, and abnormality of face, and latency to the first action potential is associated with brain atrophy. At the same time, increased soma Ca^2+^ concentration but not dendrite Ca^2+^ concentration is associated with motor or movement impairment (Fig. 6A).

**Fig. 6.**
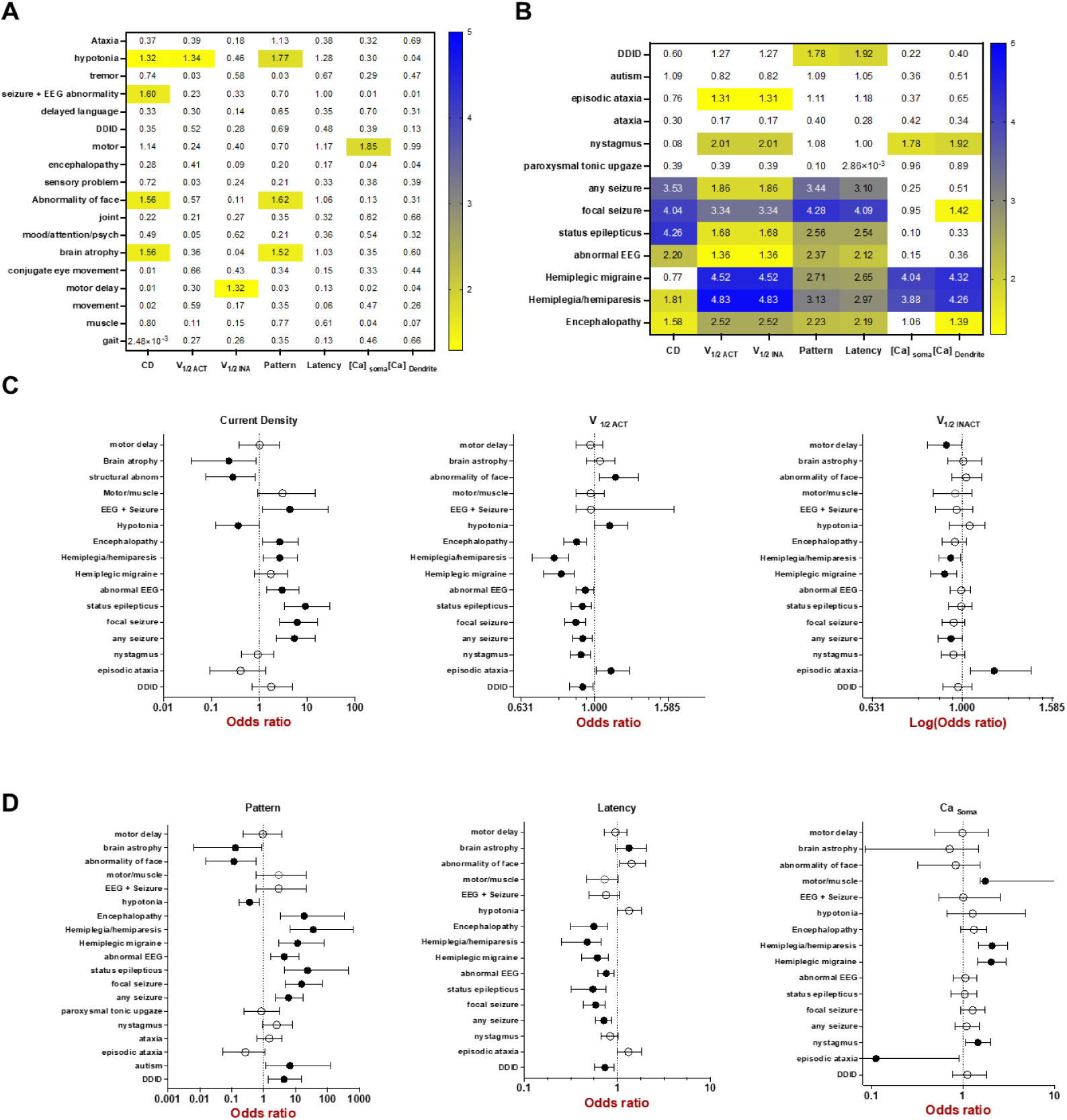
Correlations between measured and simulated neuronal function and clinical phenotypes. **A,** -log(P) values of linear regression between three measured hCa_V_2.1 properties and four simulated Purkinje function and observed 18 clinical phenotypes from 34 patients from the DD/ID cohort. **B**, -log(P) values of logistic regression analyses between molecular function and 13 clinical phenotypes in 81 patients from the review study (bottom). **C,** Forest plots of odds ratio in logistics regression analyses for current density (CD), V_1/2 ACT_ and V_1/2 INACT_ with 13 phenotypes that showed significant association with at least one measured metric; solid circle indicate significant associations; 95% Cl likelihood range for odds ratio is shown in each horizontal bars. **D,** Forrest plots of odds ratio for firing pattern, latency, Ca^2+^ concentration in soma and Ca^2+^ concentration in dendrites with 13 clinical phenotypes similarly as in **C**.

Similar analyses were performed with the clinical phenotypes in the literature review dataset, revealing that 9 (of 12) clinical phenotypes are significantly associated with at least one biophysical parameter (Fig. 6B). Strikingly, current density is significantly associated with multiple seizure phenotypes (p<0.0001), while the change in V_1/2-ACT_ is more significantly associated with hemiplegia or hemiplegic migraine (p<0.0001). Notably, V_1/2-ACT_ and V_1/2-INACT_ are both associated with episodic ataxia and nystagmus, but not ataxia. When we use the Purkinje output as the primary input, we found an additional association with DD/ID, epilepsy, encephalopathy, and hemiplegia. Both Ca^2+^ levels in soma and dendrites are strongly associated with hemiplegia (p<0.0001).

To comprehend the direction and effect size of the statistically significant associations, we plotted the odds ratios (OR) of the associations between the six clinical phenotypes from the first dataset and nine from the review study in forest plots with metrics of molecular and neuronal function (Fig. 6C-6D).

As shown in Fig. 6C, a 1 unit (1 fold from WT) increase in current density is positively associated with the risk for any seizure (OR=5.4, 95% Cl = 2.3 to 14.4, p =0.0002), focal seizure (OR=6.1, 95% CI = 2.6 to 16.3, p =9.0 × 10^-5^) and status epilepticus (OR=9.1, 95% Cl = 2.3 to 14.4, p=5.5 × 10^-5^), and to a lesser effect with hemiplegia/hemiparesis (OR=2.7, 95% Cl = 1.2 to 6.1, p=0.015). On the other hand, a 1 unit reduction (e.g., complete loss of function) in current density is significantly associated with abnormality of face (OR=4.3, 95% Cl = 1.13 to 13, p=0.044), and trending in significant association with hypotonia (OR=2.7, p=0.055) and brain atrophy (OR =3.6, β=0.058). In comparison, each mV right-shift of V_1/2-ACT_ is significantly associated with greater risk for hypotonia (OR=1.1, p=0.045) and episodic ataxia (OR = 1.1, p=0.03), while each left-shift mV of V_1/2-ACT_ is significantly associated with epilepsies with OR ranging from 1.06 to 1.13, and hemiplegia with OR = 1.30 (p= 1.5 × 10^-5^). When simulated neuronal metrics were used (Fig. 6D), we observed variants that produced tonic firing are associated with hypotonia (OR=2.8, p=0.017), brain atrophy (OR =8.3, p=0.03) and abnormality of face (OR= 9.09, p=0.024), while variants associated with complex firing are associated with epilepsies (OR between 6 and 23), and most strikingly with hemiplegia (OR = 35). Across all phenotypes in the two datasets, these associations are consistent with the idea that pathogenic variants that induce greater calcium influx are associated with seizures and hemiplegia, while LoF mutations are associated with brain atrophy, abnormality of face, hypotonia and trending in significance with episodic ataxia. Interestingly, while the results are consistent across both datasets for epilepsy or abnormal EEG, there is no significant association between any biophysical metrics and autism or DD/ID phenotype in the first dataset, while hyperpolarized V_1/2-ACT_ in the second dataset showed a significant association with DD/ID and autism. The ascertainment criteria, the number of patients in each dataset, and the nature of the variant in each dataset (∼50% of variants associated with complex firing in the first dataset, vs 75% variants in the second dataset), as well as the source of the phenotypes used may influence analyses performed.

## DISCUSSION

Pathogenic variants of *CACNA1A* have been identified in patients with hemiplegic migraine, autism, epilepsy, and ataxia(*38*). These coding variants often exert distinct and sometimes opposite effects on Ca_V_2.1 function. Therefore, the channelopathy surrounding *CACNA1A* has remained elusive. The highly diverse impact of coding mutations on molecular function of Ca_V_2.1 confounds our ability to address if and how a given allele of *CACNA1A* contributes to disease pathophysiology. In this study, we analyzed the function of an entire allelic series of *de novo* variants of *CACNA1A* in a neurodevelopmental disorder cohort, together with the most common coding variations of the same gene in the population (gnomAD), using a highly sensitive assay of channel protein function. While none of the common *CACNA1A* variants we analyzed significantly altered any of the five biophysical parameters (Fig. 1), all but one of the *de novo* changes significantly altered at least one aspect of properties that we examined. Strikingly, we found that the most common molecular phenotype induced by *de novo* missense variants of *CACNA1A* is the complete loss or significant reduction of Ca_V_2.1 current density (Fig. 1C) in our experimental setting. Furthermore, of the 22 variant channels that produced measurable currents, 15 (70%) changed how channels activate upon voltage depolarizations, and 11 (50%) altered how channels inactivate to changes in membrane potentials (Fig. 1C). Notably, a few variants significantly slowed the rate of the channel entering the inactive state or return to the closed state (from the open state) of the channel, often concurrent to the changes in voltage dependence of activation. Interestingly, most *de novo* mutations of *CACNA1A* did not recur, while only six (V176M, A713T, R1349Q, V1393M, D1634N, and R1664Q) variants appeared in more than one patient in the cohort. Among the six, V176M did not significantly alter any channel properties; three variants (A713T, R1349Q, V1393M) left-shifted the voltage dependence of activation, and were present in a total of 12 patients, while two variants (D1634N, R1664Q) encoded channels whose currents we couldn’t detect and were present in 5 patients. Taken together, our results underscore the significant impact of *de novo* missense variants of *CACNA1A* in the risk for neurodevelopmental disorders and emphasize the role of missense change in *CACNA1A* in the context of human disorders in general.

### Purkinje excitability simulation and neurodevelopmental disorder

Perhaps somewhat surprisingly, we identified variants that encode hCa_V_2.1 channels with both increased and decreased activities in the disease cohort, highlighting the vulnerability of neurodevelopment to impaired Ca_V_2.1 function, with both GOF and LOF underlying disease etiology. Some variants displayed mixed phenotypes. For example, G701E significantly reduced Ca_V_2.1 current density (LOF), with a large leftward shift in voltage dependence of activation (GOF), demonstrating complexity of interpretating channel biophysical changes in terms of GOF or LOF in the neuronal output, as we have discussed previously(*18*). To address this, we used NEURON simulations to compute the output of Purkinje neurons as an omnibus measure of Ca_V_2.1’s physiological impact, as emerging evidence suggests that altered Purkinje excitability is a major driver for motor dysfunction in a subset of ataxias that are common in *CACNA1A* channelopathy(*39*). The Purkinje neurons modulate information exchange between climbing/parallel fibers and the deep cerebellar nuclei (DCN). Cerebellum not only plays an important role in motor control and planning as previously well known, but also is increasingly recognized for its contribution to emotion processing and cognitive function(*40–43*).

Purkinje neurons rely on Ca_V_2.1 channels to regulate their intrinsic pacemaking activity, displaying irregular firing in animals with impaired hCa_V_2.1 activity(*44–47*). Using this computational model, we simulated the impact of heterozygous *CACNA1A* missense variations on Purkinje output. We found that hCa_V_2.1 haploinsufficiency produced spontaneous tonic firing similar to WT but with a much longer latency to the first action potential. Perhaps most strikingly, the left shift of V_1/2-ACT_ activation showed drastic changes to the default spontaneous tonic firing, leading to various complex bursting firing patterns that may significantly alter Purkinje neuron output (Fig. 4A, Suppl. Fig. 5).

Altered Purkinje activity may change its inhibitory input to DCN neurons, which send the major cerebellar output to other regions of brain. Interestingly, both reduced Purkinje neuron function and hypoactive DCN are observed in different forms of ataxia(*39, 45*). This is consistent with our analyses that both LOF and GOF *CACNA1A* missenses were found in patients with ataxia; however, episodic ataxia is only significantly associated with LOF variants (Fig. 6), supporting a precise molecular mechanism for specific clinical outcomes. In addition to its role in motor function, the cerebellum is increasingly recognized as a key node in neurodevelopmental disorders(*42*). Early cerebellar damage is often associated with poorer outcomes than cerebellar damage in adulthood, suggesting that the cerebellum is particularly vulnerable during development. Impaired cerebellum function early in development can have long-term effects on movement, cognition, and affective regulation. We simulated the impact of *de novo CACNA1A* missense on Purkinje neuron output and found that *CACNA1A de novo* variants greatly altered the Purkinje firing pattern, presumably leading to significant change in cerebellum output during development, supporting the hypothesis that cerebellar function may mediate *CACNA1A* channelopathy in neurodevelopmental disorders(*48*).

### Structural insights

The most significant molecular phenotypes of *de novo* coding variants of *CACNA1A* involve changes in channel current density (73% of total) and altered voltage-dependence of activation (68% of those with detectable currents). Reduction in current density may be induced by altered expression, stability, or trafficking of the channel protein or by changing the conduction of the channel pore, entailing additional biochemical and biophysical analyses to dissect the molecular mechanisms that are beyond the scope of this report. We mapped variants affecting either current density or V_1/2-ACT_ in the 3D structure of hCa_V_2.1 channel (Fig. 3B-3D), and they are distributed across the pore domain (PD) and transmembrane voltage sensing domain (VSD) of the hCa_V_2.1 channel. Three regions are highlighted in hCa_V_2.1 that contained amino acid changes that influenced the channel currents (Fig. 3C). The S5 and S6 segments of the PD, along with the S4 segments of the VSD and S4-S5 linker, harbored the majority of the *de novo* variants that produced hCa_V_2.1 channel with significantly reduced current density, suggesting that amino acid changes in these regions may induce change in channel protein production, stability, trafficking, or pore conduction, all of which may lead to reduced total current density. Missense changes that alter the V_1/2_ activation are also mapped to PD and VSD domains in hCa_V_2.1 (Fig. 3D). Perhaps expectedly, two arginine residue changes (R1349Q and R1352Q) induced substantial shifts in the V_1/2_ activation due to their role in voltage sensing in the S4 voltage sensing segment. Somewhat surprisingly, right shifts induced by I1357M and F1367I (19 and 27 mV in ΔV_1/2-ACT_ compared to WT) are located in the S4-S5 linker, where the voltage-sensing is transduced to the opening of the pore. I1357M and F1367I may impede such transduction and make the channel harder to open. Overall, our findings align with previous known mutations in the VSD and PD regions of voltage-gated calcium and sodium channels(*49–51*), where mutations in these domains alter current density and voltage-dependence of activation. In addition, our results revealed many 3D structural mapped missense changes in *CACNA1A* that significantly altered the channel function, offering fine-tuned metrics on developing structure-based predictions of *CACNA1A* variants that may affect voltage-dependent activation and current density. In the long term, functional studies at scale, coupled with insights from 3D structural information, will train future algorithms for more precise functional prediction in voltage gated ion channels, facilitating more precise genetic - molecular - clinical phenotype analyses.

### Disease associations and molecular/clinical phenotypes correlations

Most genetic analyses from large cohort studies focus on protein truncation variants because existing algorithms often struggle to differentiate benign coding variants from those with functional impact(*52–55*). Using our functional data, we evaluated the predictions of AlphaMissense (AM) against all missense variants in gnomAD (Fig. 5) and found that it correctly predicted functional impacts with more than 85% and 95% accuracy for gnomAD and de novo coding variants, respectively. This lends us confidence in using AlphaMissense to predict the pathogenicity of coding variants of *CACNA1A* in epilepsy cohorts (Fig. 5C-5D). Our analyses revealed that pathogenic missense variants of *CACNA1A* are a significant contributor to epilepsy in general and greatly increase the risk of developmental and epileptic encephalopathy (DEE) in particular (Fig. 5C), consistent with previous anecdotal reports with *CACNA1A* mutations(*16, 56, 57*). However, as shown in Fig. 5D, while AlphaMissense accurately predicted the pathogenicity of *CACNA1A* variants in our analyses, it performed poorly when predicting the distinct biophysical impact or the direction of the impact (LOF or GOF) of individual variants, substantiating the need for precise functional studies in channelopathy until we have better predictors.

Using precise functional measurements of hCa_V_2.1 variant channel properties, along with two datasets of clinical phenotypes, we found that increased current density and left shift in V_1/2_ activation are significantly associated with focal seizure, status epilepticus, EEG abnormalities, hemiplegic migraine, or hemiplegia (Fig. 6A), supporting that greater calcium influx mediated by Ca_V_2.1 channels underlie epileptic phenotypes as well as hemiplegia. In contrast, LOF mutations are significantly associated with hypotonia, episodic ataxia, brain atrophy, and abnormality of face, suggesting distinct aspects of Ca_V_2.1 function may drive the different facets of disease pathophysiology. Depending on the completeness and accuracy of clinical descriptions, together with temporal pattern of initial clinical features, these analyses start to reveal the impact of molecular function on clinical phenotypes measured in individual patients and help in the development of precision treatments for the spectrum of disorders caused by *CACNA1A* dysfunction.

## Conclusions, limitations, and future directions

This is a comprehensive study with consistently and similarly measured molecular function of an entire allelic series of *CACNA1A* missense variants derived from a cohort with annotated clinical phenotypes to address the potential relationships between molecular and clinical phenotypes. This study focused on biophysical properties of hCa_V_2.1 using the heterologous expression system in the presence of auxiliary subunit β_3_ and did not address whether the impact of the missense changes may be different in the presence of a different β subunit or together with auxiliary subunit α2δ. However, we did evaluate the impact of two variants (L618S and C1369R) in the presence of β_4_, and the results are similar to the data presented this report (ΔV_1/2-ACT_ = - 7.3 ± 0.8 mV, p<0.001 and ΔV_1/2-ACT_ = - 13.9 ± 1.6 mV, p<0.001 in β_4_, while they were ΔV_1/2-ACT_ = - 7.7 ± 0.9 mV, p<0.001 and ΔV_1/2-ACT_ = - 15.6 ± 0.7 mV, p<0.001 in β_3_ for L618S and C1369R, respectively), validating our general approach on delineating biophysical impact of the missense changes. Still, future studies including different auxiliary subunits and neuronal context may reveal additional neuronal impact of these missense changes and shed light on neuronal mechanism of the deficits. Of note, 14 (of 34, 40%) patients in clinical dataset one and 72 (of 81, 88%) patients in clinical dataset two have recurring mutations. In comparison, only 18 (of 58, 30%) patients have recuring *CACNA1A de novo* changes in the 31,000 trio-cohort. Hence, our molecular function – clinical phenotype association analyses may have been biased towards recuring mutations. Nevertheless, with the expanding and increasingly comprehensive clinical phenotyping of patients, along with detailed molecular function annotation on missense changes, this approach will ultimately address whether different Ca_V_2.1 dysfunction underlie similar, distinct or overlapping clinical conditions to guide treatment selections. Given that both GOF and LOF of *CACNA1A* are found in patients of neurodevelopmental disorder, understanding the underlying neurobiological basis of the disease mechanism and progression, and defining the normal range of Ca_V_2.1 activity, will be the next step in designing the intervention strategies for specific mutations in the future. In this regard, our modeling of Purkinje excitability suggests distinct patterns of Purkinje firing (tonic and complex bursting) may underlie disease pathophysiology, consistent with recent literatures highlighting the cerebellum’s function in normal and pathological neurodevelopment.

## MATERIALS AND METHODS

### Molecular and cell biology Reagents

The reference CACNA1A cDNA construct (NM_001127221.2; NP_001120693.1) was inserted into the pIR-CMV-IRES-mScarlet) expression vector (Addgene plasmid # 162279). To generate 50 coding cDNA variants, site-directed mutagenesis was performed, and mutated sites were validated by Sanger sequencing. The HEK293 cell line that expresses the Ca_V_ auxiliary β3 subunit (a gift from Diane Lipscombe) was constructed using the T-ReX system (Thermal Fisher). The Ca_V_ β3 subunit is one of the four auxiliary Ca_V_ -β subunits that contain interaction domains that are responsible to traffic Ca_V_ 1 and Ca_V_ 2 family channels to the plasma surface. HEK293 cells stably expressing the Ca_V_β_3_ subunit were grown in DMEM/F12 +GlutaMAX media (Thermo cat. 10565018) containing 10% FBS (Gibco, cat. 26140079). The expression of Ca_V_β_3_ was induced by 1 μg/ml doxycycline (Thermo cat. J60579-22). The detailed culturing conditions were previously described(*18*).

### Transient transfection for electrophysiology

Electroporation was performed with the MaxCyte ATx^TM^ system. Cells were seeded at a density of 1.5-2 million cells per T175 flask in 30mL of media supplemented with 30µL of 1 mg/mL doxycycline, three to four days before the electroporation. On the day of electroporation, cells of 70-85% confluent were washed with 5mL of PBS once and incubated in 3mL of dissociation solution (MaxCyte, TrypLE) at room temperature (RT) until cells were detached, followed by adding 7 mL of DMEM/F12 + 10% FBS quenching media (Thermo cat. 10565018). Cells from multiple T175 flasks were pooled into a 50 ml conical tube and centrifuged at 1000rpm for 5 minutes at RT. The supernatant was aspirated, and washed in 2 ml of electroporation buffer (MaxCyte, EPB-1) and subsequently centrifuged at 1000 rpm for 2 minutes. The supernatant was aspirated, and then the pellet was resuspended in the electroporation buffer at a concentration of 120 million cells per ml in a 1.5 ml microfuge tube. 20 µg (5ug/µl) of the CACNA1A plasmid encoding WT or a variant channel was aliquoted into a microfuge tube with HyClone electroporation buffer (Cytiva cat: EPB1) to the final volume of 13 µl. An aliquot (87µL) of the cell suspension was then mixed with the premade 13 µl DNA solution containing 20µg of CACNA1A DNA and pipetted gently into the OC100-2X cassette. Ten million cells were used for each electroporation using the standard HEK293 protocol. After electroporation, 10µl DNase I (Cat#07900, STEMCELL Technology, 1ug/ul) was added to the cuvette, and the electroporated cells were then gently pipetted into a tissue culture dish, placed into a 37℃ incubator for 30 minutes, re-suspended in DMEM/F12 media supplemented with 10% FBS, then replated onto 10 cm dishes in the presence of doxycycline. After a 48-hour incubation at 37℃, cells were harvested for electrophysiology experiments. The overall percentage of detected currents across all variants and controls is shown in Suppl. Fig. 1

### Electrophysiology Solution

All recording solutions were prepared with ultrapure Milli-Q water (18 MΩ-cm), filtered with 0.22 µm PES membrane, and stored at 4°C until use. External recording solution was (in mM): 10 HEPES, 142 TEA-Cl, 5 D-Glucose, 5 BaCl_2_, 300 mOsm, and pH 7.4 (TEA-OH). Internal recording solution was (in mM): 10 EGTA, 10 HEPES, 90 Cs_2_SO_4_, 1 MgCl_2_, 17 CsCl, 290 mOsm pH 7.2 (CsOH). The remaining solutions, including those used to promote the electrical seal formation and other technical details, were published previously(*18, 20*). All salts were purchased from Sigma-Aldrich. Hygroscopic reagents were kept in a salt desiccator.

### Automated patch-clamp experiments

All electrophysiology experiments were performed using the SyncroPatch 384PE platform (Nanion Technologies®), as described previously(*19*) In brief, cells were harvested 48 hours after electroporation, and the experiments were performed within one hour. The assays were conducted in single-hole chips with aperture resistances of 4-5 MΩ. Before addition of the cells into the chamber, the junction potential (∼12 mV) and the fast-capacitive component were compensated once in whole-cell configuration, the slow capacitive component (C_slow_) was canceled, and the series resistance (R_s_) compensation was set to 80%. For each 384-well plate, cells expressing the WT hCa_V_2.1 were always used as the control, and the analyzed properties of each variant channel are compared to those of the WT hCa_V_2.1 control in the same plate. WT and hCaV2.1 channels were randomly positioned on the 384-well plate to prevent any issues caused by their location. We typically recorded 5 variant channels in each 384-well plates in parallel with the WT channel. The data were acquired at 20 kHz and filtered at 10 kHz using Nanion Technologies® proprietary software Patch Control 384 (V.1.4.5). The data were exported and processed with DataControl384 (Nanion) for further offline biophysical analyses. In the absence of b3 induction, there is little expression of hCa_V_2.1 currents (Suppl. Fig. 3), and in the presence of WT transient transfection, the average of the current density, across multiple batches, was ∼22.2± 10.7 pA/pF.

### Electrophysiology

#### Electrophysiological protocols and biophysical analyses

The holding potential was set to −90 mV for all voltage protocols. Peak current and the activation steady-state parameters were obtained from Ca_V_2.1 currents elicited by 250 ms depolarization steps from −50 to +50 mV at 5 mV increments per sweep, followed by a 250 ms test pulse at −35 mV. The current-voltage relationship (*I-V*) was constructed by plotting the maximum peak current magnitude, normalized by the cell capacitance, as a function of the voltage applied. The voltage-conductance relationship (*G(V)*) was calculated by dividing the maximum current magnitude at each voltage by the corresponding driving force:

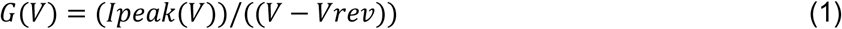

The reversal potential (*V_rev_*) was calculated from the intersection with the X-axis of the linear extrapolation of the last four points of the I-V curve. The voltage-dependent activation parameters were obtained by fitting a single Boltzmann function to the normalized conductance (*G(V)/Gmax*):

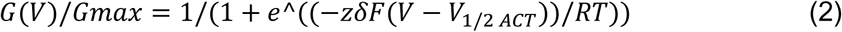

*V_1/2-ACT_* is the voltage at which half of *Gmax* is reached; *zδ* corresponds to the slope of the Boltzmann function (voltage sensitivity); *R* corresponds to the universal gas constant; *T* is the absolute temperature in Kelvin; and *F* is the Faraday constant. The mean midpoint values were represented as described by Horrigan and Aldrich(*21*).

Steady-state inactivation parameters were obtained by a 5 s prepulse from −100 mV to +30 mV at 10 mV increments per sweep, followed by a 250 ms test pulse at 15 mV (Suppl. Fig. 2). The current amplitudes (*I*) at the test pulse were normalized to the maximum current amplitude at −100 mV (*Imax*) and represent the fraction of channels that were able to open at a given prepulse. The voltage-dependent inactivation relationship was fitted by a Boltzmann function according to the following equation:

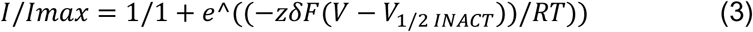

where *V_1/2-INACT_* is the voltage at which half of *Imax* is reached; *zδ* corresponds to the slope of the fitted Boltzmann function (voltage sensitivity); and *R, T, and F are the same as in equation 2*.

The tail current was obtained by applying a voltage protocol that includes −90 mV holding potential, +25 mV (test pulse) for 50 ms, and voltage steps (Δ10 mV) from −80 mV up to +20 mV (pre-pulse) for 250 ms. A depolarizing pulse to +25 mV opens channels, and the following step pulse repolarizes the membrane potential before the inactivation process starts. The time constant associated with the closing of the channel (τ_DEACT_) is described by an exponential decay function fitted to the tail protocol.

Window current was determined using the following equation: (1/(1+exp((*V*-*V_1/2 ACT_*)/*k_ACT_*)) x ((1-C)/(1+exp((*V*-*V_1/2 INACT_*)/*k_INACT_*)) + C), where *k_ACT_* is the slope factor of activation, *V* is the test potential, and *k_INACT_* is the slope factor of inactivation, and C is constant. It was obtained from the collected data of WT and variants using programs written in Igor Pro 6 (Wavematrics, Lake Oswego, OR).

#### Biophysical parameter normalization

To calculate the change in the midpoint of the activation and inactivation voltages (ΔV_1/2-ACT_ and ΔV_1/2-INACT_) for the analyzed Ca_V_2.1 channels, the values for each cell expressing wild-type (WT) hCa_V_2.1 (equation 4) or a variant channel (equation 5) were normalized to the mean WT hCa_V_2.1 values for V_1/2-ACT_ and V_1/2-INACT_ and presented as the mean shift in mV from the mean value of WT hCa_V_2.1 channels.

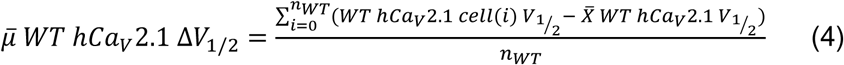

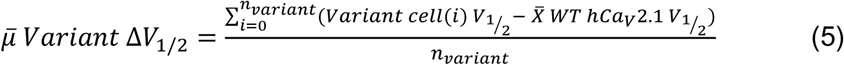

The peak current density and recovery time from inactivation of the variant channels are presented as the fold change with respect to the mean values of WT hCaV2.1 channels. We used equation 6 for WT and equation 7 for variant channels.

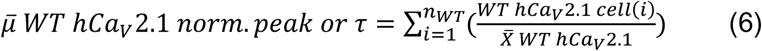

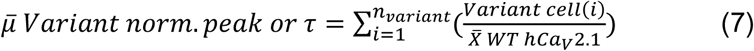

To calculate the z-scores, the specific biophysical parameters (voltage dependence activation and inactivation, current density, recovery from inactivation) of the different variants were transformed into dimensionless distance from the WT properties measured by the standard deviation according to the following equation:

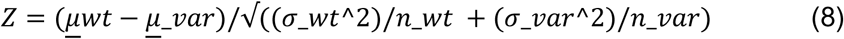

where *μ*, *σ^2^*and *n* stand for the mean, standard deviation, and size of the sample, respectively.

### Statistics

For the electrophysiology recordings, in total, we analyzed 419 cells that expressed WT Ca_V_2.1 channels and 839 cells that each expressed one of the 50 variants of Ca_V_2.1 channels. Each hCa_V_2.1 variant channel was recorded from at least ten cells. The quality control parameters for the SyncroPatch recording and analyses were previously described(*20*). Fifteen batches of experiments were run to characterize 50 *CACNA1A* variants. We recorded five variant channels and the WT channel on each 384-well plate. Every parameter obtained for each cell carrying a variant was normalized to the mean WT hCa_V_2.1 signal from the same plate. Then, we performed ANOVA for the normalized values of each variant and the normalized values for the WT channel within the same batch, followed by Dunnett’s post hoc correction for multiple comparisons. The significance threshold was defined as *p* ≤ 0.05 after the post hoc correction. Data are presented as the mean ± S.E.M., unless stated otherwise. For other analyses, ANOVA was performed for group comparisons followed by Dunnett’s post hoc comparison unless otherwise noted.

### Structural model of hCa_V_2.1

The apo structure of human Ca_V_2.1 (A0A1B0GUS3_HUMAN) channel was obtained from the Alphafold2 database. Next, unresolved cytosolic regions and linkers between repeats in the template were predicted using I-TASSER(*17*) to connect the transmembrane repeats. Subsequently, the final model underwent refinement by selecting lists of 3D predictions for each unresolved region, guided by secondary structure prediction, including Jpred(*22*).

### *NEURON* Simulations

To predict the impact of Ca_V_2.1 variants on neural activity, we employed a comprehensive multi-compartment biophysical model of the Purkinje neuron(*23*) (ModelDB, 229585). This model, implemented in NEURON, encompasses ionic conductance across somatic, dendritic, and axonal compartments, providing a detailed morpho-electrical representation of the Purkinje cell. Its architecture includes one soma, a few proximal dendrites, over 1500 distal dendrites, and a detailed axonal geometry. Voltage- and Ca^2+^-dependent mechanisms are distributed among these compartments. To evaluate Purkinje excitability, we simulated the model for 5 seconds from identical initial conditions with different Ca_V_2.1 biophysical properties, recording membrane voltage on an axon collateral and dendritic calcium on a distal dendrite. Steady-state calcium concentrations in the soma and distal dendrites, as well as the firing pattern derived from the voltage trajectory measured on the axon, were assessed. To mimic heterozygosity, 50% of simulated Ca_V_2.1 channels were loaded with specific variant biophysical properties, while the remaining 50% followed wild-type (WT) parameters. From the simulations for each variant channel, we obtained five derived parameters: pattern (tonic =0, complex bursting =1), latency (milliseconds to the first action potentials), rate (average action potential per second during the simulated 5 seconds), Ca soma concentration ( steady state calcium concentration between 2.5 s and 5s in the cell soma in mM), Ca dendrite concentration (steady state Ca concentration in mM between 2.5 and 5 seconds in the simulation).

### AlphaMissense prediction and epilepsy cohort analyses

We downloaded the AlphaMissense predictions and analyzed the pathogenicity scores of missense variants in *CACNA1A* discovered in different disease cohorts and the general population. This included 42 *de novo* missense variants found in individuals with neurodevelopmental disorders, 201 rare variants in individuals with epilepsy across three subtypes (36 in developmental and epileptic encephalopathy [DEE], 71 in genetic generalized epilepsy [GGE], and 94 in non-acquired focal epilepsy [NAFE]) as well as 250 in controls sequenced in the same epilepsy WES study(*24*). Additionally, we examined 659 missense variants reported in gnomAD(*25*) across the entire allele frequency spectrum. Of the 50 variants that we characterized, I346Del and K1202Del could not be assessed using AlphaMissense.

### Clinical Phenotypes

We obtained standardized clinical terms for 34 individuals (with 18 unique variants) that carried *de novo* missense variants in *CACNA1A* from GeneDx LLC. Data were provided under a Data Transfer Agreement between the GeneDx Corporation and the Broad Institute. Phenotypic terms were standardized to the Human Phenotype Ontology (HPO; hpo.jax.org). Among the 18 unique *de novo* variants identified, six occurred more than once, with A713T (present in 6 individuals) being the most frequent. We extracted the most frequent 18 clinically relevant phenotypic labels from the HPO terms from this dataset. In parallel, we queried a dataset constructed from individuals reported in the literature (397) and clinical cases at three institutions (57), totaling 454 individuals with *CACNA1A-*related conditions, including 301 total unique *CACNA1A* variants. Here, we identified 81 patients carrying 22 unique missense *CACNA1A* variants and extracted 13 clinical features(*26*). Thirteen of the 22 variants were recurrent missense changes, with the most frequent being R1352Q, present in 22 patients. We then correlated the incidence of these clinical features with the measured and simulated electrophysiology parameters associated with each variant. Logistics regressions were used to correlate the molecular phenotypes with the clinical phenotypes.

## Abbreviations

LoF: loss-of-function
GoF: gain-of-function

## Data availability

Raw data were generated at the Stanley Center for Psychiatric Research at the Broad Institute. Derived data supporting the findings of this study are available from the corresponding author on request.

## Supplementary materials

This Manuscript file includes:

Material and Methods

Suppl. Fig.1 to Fig.5

Suppl. Table 1 to Table 3

## Acknowledgments

The authors thank Dr. Lei A Wang for the initial modeling in Purkinje Neurons. We thank Dr. Dennis Lal for very helpful discussions during the study. We thank the CACNA1A Foundation for the inspiration to find a cure for patients with CACNA1A mutations. We thank Young Chueng and Maofu Liao for providing the homology model of Ca_V_2.1.

## Authors’ Contributions

JQP conceived and supervised the project. JQP wrote the manuscript together with EK, SC, DB, and NB. NB, SJ, and EK performed the Syncropatch experiments. EO performed NEURON simulations together with DB. EK performed structural analyses and analyses with clinical phenotypes. SC and MD calculated the pathogenicity score for missense variants using AlphaMissense and performed the genetic analyses in epilepsy cohorts. AP, LS, LL, IH provided the additional *de novo* variants to be included in the analyses and the discussion of clinical phenotypes. SM and AL provided clinical phenotypes in the cohort study and LL and IH in the review study. AG provided the plasmid backbone, and provided valuable feedback during the project.

