## Supplementary material for "Characterization of the functional and clinical impacts of CACNA1A missense variants found in neurodevelopmental disorders": Supp. Table 1

| | | Current density | V1/2_act | V1/2_inact | tau_inact | tau_deact |
|---|---|---|---|---|---|---|
| | | z-score | z-score | z-score | z-score | z-score |
| | WT | 0 | 0 | 0 | 0 | 0 |
| DI | V176M | -0.6804297 | -0.8111481 | -0.0803895 | 1.8352516 | -0.3423517 |
| | A222V | -5.0031941 | -1.0992257 | -1.2370026 | 0.59821321 | 2.3314162 |
| | P269L | -2.9258465 | -1.8513722 | 2.98110914 | -0.0650682 | 1.05976155 |
| | I345M | -4.1544719 | -1.4986711 | -2.5925107 | -0.6823997 | -0.1295436 |
| | WT | 0 | 0 | 0 | 0 | 0 |
| DII | R481G | -0.7077999 | -2.8843028 | 3.11901569 | 0.78135411 | -0.9598836 |
| | L618S | 2.81197042 | 6.79356356 | 3.33831782 | 2.9194479 | 4.43418306 |
| | G701E | -7.8204228 | 12.6152661 | 0.16718099 | 0.55398343 | 2.92345394 |
| | A713T | 1.82448603 | 9.37296424 | 0.70022253 | 3.61335279 | 1.24650291 |
| | V714M | -4.7757431 | -0.4462136 | -1.7072659 | -0.21367 | 0.297609 |
| | WT | 0 | 0 | 0 | 0 | 0 |
| DIII | I1339S | -4.5363448 | 3.8649339 | -0.950023 | -0.705372 | 1.1030765 |
| | R1349Q | -2.4921173 | 10.0900264 | -2.3061344 | 0.34849452 | -0.9340392 |
| | R1352Q | -2.3305398 | 6.50140069 | -4.2014376 | -0.6469768 | -0.1580309 |
| | P1353L | -6.268822 | 2.83439644 | -2.2784148 | 1.21862 | 2.8990967 |
| | I1357M | -3.6289135 | -8.4255641 | 4.32018813 | 0.50755892 | 0.50057446 |
| | R1359W | -2.7029756 | -0.4612292 | 0.55556614 | -0.9373913 | 2.74086543 |
| | F1367I | -0.9724245 | -17.777592 | 3.26615091 | 0.5077346 | -0.6219357 |
| | C1369R | 2.71927969 | 5.1250706 | -1.3089366 | 1.00955217 | 0.42595873 |
| | V1377I | -8.6175275 | -6.307966 | 2.25329025 | -0.4862354 | 4.01808819 |
| | V1393M | 6.40023181 | 2.74762548 | -6.7465903 | -1.4442009 | 4.52140083 |
| | V1393L | 3.43309928 | 11.4430588 | -4.429957 | -2.8224947 | 2.53098893 |
| | A1508T | 1.45717755 | 11.1496671 | 0.49883561 | 3.32793647 | 2.44950778 |
| | WT | 0 | 0 | 0 | 0 | 0 |
| DIV | I1709T | 0.63991772 | 4.12194788 | -0.3671117 | 0.30344262 | 0.02429074 |
| | WT | 0 | 0 | 0 | 0 | 0 |
| gnomAD | A453T | -0.6367157 | 0.23202941 | -0.218618 | 0.60922675 | 0.85526505 |
| | E731A | -0.3530906 | -0.452717 | -0.1947701 | 0.88336288 | -0.9554856 |
| | E917D | -0.7316565 | 0.41670508 | 0.21239121 | -0.1786854 | -0.3682811 |
| | E992V | -0.3526898 | -0.1021685 | -0.4877094 | 0.65253506 | -0.9025618 |
| | P1010A | -0.408252 | -0.7769301 | -0.1004728 | 1.34643184 | -0.1324604 |
| | E1014K | -0.5086643 | -0.5809841 | 0.56344936 | -0.8877893 | -0.1470517 |
| | G1104S | -0.5128895 | -0.4060997 | 0.53312573 | 1.50906813 | -0.2758714 |
| | K1202Del | 0.51632483 | -0.6595317 | 0.2377394 | 1.06492369 | -0.459586 |
