## Supplementary material for "Characterization of the functional and clinical impacts of CACNA1A missense variants found in neurodevelopmental disorders": Supp. Table 2

| Variant | Pattern | Latency (ms) | Rate (spk/s) | Ca+2 soma (uM) | Ca+2 dend (uM) |
|---|---|---|---|---|---|
| WT | Tonic | 288.4 | 54.3333333 | 0.06838509 | 0.00833026 |
| LOF | Tonic | 532.55 | 50 | 0.04727939 | 0.00438705 |
| V176M | Tonic | 391.675 | 50.3333333 | 0.06107881 | 0.00581088 |
| A222V | Tonic | 492.475 | 50.6666667 | 0.05063603 | 0.00467218 |
| P269L | Tonic | 411.7 | 51 | 0.05845915 | 0.00554459 |
| I345M | Tonic | 389.75 | 52.3333333 | 0.05662836 | 0.00576426 |
| R481G | Tonic | 209.3 | 64 | 0.07550102 | 0.01353155 |
| L618S | Complex | 68.825 | 14 | 0.05162449 | 0.4064157 |
| G701E | Complex | 30.95 | 152.666667 | 0.200372 | 4.46760644 |
| A713T | Complex | 31.825 | 10 | 0.06955507 | 0.86398276 |
| V714M | Tonic | 527.6 | 50 | 0.04895745 | 0.0044227 |
| R1349Q | Complex | 5.75 | 164.666667 | 0.46294679 | 7.96841953 |
| R1352Q | Complex | 46.25 | 150.333333 | 0.16016526 | 3.94302054 |
| P1353L | Tonic | 354.025 | 54.3333333 | 0.05505313 | 0.00632617 |
| I1357M | Tonic | 532.125 | 49.3333333 | 0.05065155 | 0.00439531 |
| R1359W | Tonic | 494.325 | 50.6666667 | 0.04996689 | 0.00465963 |
| F1367I | Tonic | 532.525 | 49.3333333 | 0.04958681 | 0.00438658 |
| C1369R | Complex | 10.5 | 58 | 0.23034774 | 4.33693855 |
| V1377I | Tonic | 528.3 | 49.6666667 | 0.0502377 | 0.00442494 |
| V1393M | Complex | 45.975 | 12 | 0.05321636 | 0.53072017 |
| V1393L | Complex | 43.7 | 12 | 0.05610007 | 0.61978978 |
| A1508T | Complex | 3.9 | 80 | 0.40380297 | 8.58236542 |
| I1709T | Complex | 22.425 | 31 | 0.12640972 | 1.96676316 |
| I1339S | Complex | 57.525 | 149 | 0.11916518 | 2.80371625 |
