## Supplementary material for "Characterization of the functional and clinical impacts of CACNA1A missense variants found in neurodevelopmental disorders": Supp. Table 3

| N | Clinical Phenotypes repeated |
|---|---|
| 1 | Abnomality of eye |
| 2 | Abnoramlity of foot |
| 3 | Abnormaility of hairline |
| 4 | Abnormal Conjugate eye movement |
| 5 | Abnormal EKG |
| 6 | Abnormal Gait |
| 7 | Abnormal behavior |
| 8 | Abnormal cardiovascular system |
| 9 | Abnormal conjugate eye movement |
| 10 | Abnormal eyebrow morphology |
| 11 | Abnormal facial shape |
| 12 | Abnormal gait |
| 13 | Abnormal gross motor development |
| 14 | Abnormal movement |
| 15 | Abnormal muscle tone |
| 16 | Abnormal of conjugate eye movement |
| 17 | Abnormal of muscle tone |
| 18 | Abnormal respiratory system |
| 19 | Abnormal vertebral morphology |
| 20 | Abnormality conjugate eye movement |
| 21 | Abnormality of Cardiovascular system |
| 22 | Abnormality of Vision |
| 23 | Abnormality of brain |
| 24 | Abnormality of cardiovascular system |
| 25 | Abnormality of conjugate eye movement |
| 26 | Abnormality of dentition |
| 27 | Abnormality of ear |
| 28 | Abnormality of eye |
| 29 | Abnormality of eye movement |
| 30 | Abnormality of face |
| 31 | Abnormality of finger |
| 32 | Abnormality of gait |
| 33 | Abnormality of heart |
| 34 | Abnormality of inner ear |
| 35 | Abnormality of jaw |
| 36 | Abnormality of liver |
| 37 | Abnormality of lung |
| 38 | Abnormality of movement |
| 39 | Abnormality of muscle |
| 40 | Abnormality of nervous system |
| 41 | Abnormality of nose |
| 42 | Abnormality of respiratory system |
| 43 | Abnormality of skin |
| 44 | Abnormality of skin pigmentation |
| 45 | Abnormality of temperature regulation |
| 46 | Abnormality of the dentition |
| 47 | Abnormality of the mitochondrion |
| 48 | Abnormality of the tongue |
| 49 | Abnormaliy of toe |
| 50 | Anxiety |
| 51 | Ataxia |
| 52 | Autism |
| 53 | Autistic behavior |
| 54 | Behavioral abnormalities |
| 55 | Brain atrophy |
| 56 | Brain hypoplasia |
| 57 | Braint atrophy |
| 58 | Delayed fine motor development |
| 59 | Delayed gross motor coordination |
| 60 | Delayed gross motor development |
| 61 | Delayed speech and language development |
| 62 | Depression |
| 63 | Developmental regression |
| 64 | Dystonia |
| 65 | EEG abnormality |
| 66 | Encephalopathy |
| 67 | Feeding difficulties |
| 68 | Flexion Contracture |
| 69 | Gait Disturbance |
| 70 | Generalized hypotonia |
| 71 | Global developmental delay |
| 72 | Growth delay |
| 73 | Hearing impairment |
| 74 | Hypertonia |
| 75 | Hypotonia |
| 76 | Intellectual disability |
| 77 | Joint hypermobility |
| 78 | Joint laxity |
| 79 | Laryngeal web |
| 80 | Macrocephaly |
| 81 | Microcephaly |
| 82 | Motor delay |
| 83 | Muscle weakness |
| 84 | Poor growth |
| 85 | Recurrent otitis media |
| 86 | Scoliosis |
| 87 | Seizure |
| 88 | Short attention span |
| 89 | Sleep disturbance |
| 90 | Stroke |
| 91 | Tremor |
| 92 | Visual impairment |

| N | Clinical Phenotypes non-repeated 1 |
|---|---|
| 1 | Abnomality of eye |
| 2 | Abnoramlity of foot |
| 3 | Abnormaility of hairline |
| 4 | Abnormal Conjugate eye movement |
| 5 | Abnormal EKG |
| 6 | Abnormal Gait |
| 7 | Abnormal behavior |
| 8 | Abnormal cardiovascular system |
| 9 | Abnormal eyebrow morphology |
| 10 | Abnormal facial shape |
| 11 | Abnormal gross motor development |
| 12 | Abnormal movement |
| 13 | Abnormal muscle tone |
| 14 | Abnormal respiratory system |
| 15 | Abnormal vertebral morphology |
| 16 | Abnormality of Vision |
| 17 | Abnormality of brain |
| 18 | Abnormality of dentition |
| 19 | Abnormality of ear |
| 20 | Abnormality of eye movement |
| 21 | Abnormality of face |
| 22 | Abnormality of finger |
| 23 | Abnormality of heart |
| 24 | Abnormality of inner ear |
| 25 | Abnormality of jaw |
| 26 | Abnormality of liver |
| 27 | Abnormality of lung |
| 28 | Abnormality of movement |
| 29 | Abnormality of muscle |
| 30 | Abnormality of nervous system |
| 31 | Abnormality of nose |
| 32 | Abnormality of respiratory system |
| 33 | Abnormality of skin |
| 34 | Abnormality of skin pigmentation |
| 35 | Abnormality of temperature regulation |
| 36 | Abnormality of the dentition |
| 37 | Abnormality of the mitochondrion |
| 38 | Abnormality of the tongue |
| 39 | Abnormaliy of toe |
| 40 | Anxiety |
| 41 | Ataxia |
| 42 | Autism |
| 43 | Autistic behavior |
| 44 | Behavioral abnormalities |
| 45 | Brain atrophy |
| 46 | Brain hypoplasia |
| 47 | Delayed fine motor development |
| 48 | Delayed gross motor coordination |
| 49 | Delayed gross motor development |
| 50 | Delayed speech and language development |
| 51 | Depression |
| 52 | Developmental regression |
| 53 | Dystonia |
| 54 | EEG abnormality |
| 55 | Encephalopathy |
| 56 | Feeding difficulties |
| 57 | Flexion Contracture |
| 58 | Gait Disturbance |
| 59 | Generalized hypotonia |
| 60 | Global developmental delay |
| 61 | Growth delay |
| 62 | Hearing impairment |
| 63 | Hypertonia |
| 64 | Hypotonia |
| 65 | Intellectual disability |
| 66 | Joint hypermobility |
| 67 | Joint laxity |
| 68 | Laryngeal web |
| 69 | Macrocephaly |
| 70 | Microcephaly |
| 71 | Motor delay |
| 72 | Muscle weakness |
| 73 | Poor growth |
| 74 | Recurrent otitis media |
| 75 | Scoliosis |
| 76 | Seizure |
| 77 | Short attention span |
| 78 | Sleep disturbance |
| 79 | Stroke |
| 80 | Tremor |
| 81 | Visual impairment |

| N | Clinical Phenotypes non-repeated 2 |
|---|---|
| 1 | Abnomality of eye |
| 2 | Abnoramlity of foot |
| 3 | Abnormaility of hairline |
| 4 | Abnormal Conjugate eye movement |
| 5 | Abnormal EKG |
| 6 | Abnormal Gait |
| 7 | Abnormal cardiovascular system |
| 8 | Abnormal eyebrow morphology |
| 9 | Abnormal facial shape |
| 10 | Abnormal gross motor development |
| 11 | Abnormal muscle tone |
| 12 | Abnormal respiratory system |
| 13 | Abnormal vertebral morphology |
| 14 | Abnormality of Vision |
| 15 | Abnormality of brain |
| 16 | Abnormality of dentition |
| 17 | Abnormality of ear |
| 18 | Abnormality of eye movement |
| 19 | Abnormality of face |
| 20 | Abnormality of finger |
| 21 | Abnormality of heart |
| 22 | Abnormality of inner ear |
| 23 | Abnormality of jaw |
| 24 | Abnormality of liver |
| 25 | Abnormality of lung |
| 26 | Abnormality of movement |
| 27 | Abnormality of muscle |
| 28 | Abnormality of nervous system |
| 29 | Abnormality of nose |
| 30 | Abnormality of respiratory system |
| 31 | Abnormality of skin |
| 32 | Abnormality of skin pigmentation |
| 33 | Abnormality of temperature regulation |
| 34 | Abnormality of the dentition |
| 35 | Abnormality of the mitochondrion |
| 36 | Abnormality of the tongue |
| 37 | Abnormaliy of toe |
| 38 | Anxiety |
| 39 | Ataxia |
| 40 | Autism |
| 41 | Autistic behavior |
| 42 | Behavioral abnormalities |
| 43 | Brain atrophy |
| 44 | Brain hypoplasia |
| 45 | Delayed fine motor development |
| 46 | Delayed gross motor coordination |
| 47 | Delayed gross motor development |
| 48 | Delayed speech and language development |
| 49 | Depression |
| 50 | Developmental regression |
| 51 | Dystonia |
| 52 | EEG abnormality |
| 53 | Encephalopathy |
| 54 | Feeding difficulties |
| 55 | Flexion Contracture |
| 56 | Gait Disturbance |
| 57 | Generalized hypotonia |
| 58 | Global developmental delay |
| 59 | Growth delay |
| 60 | Hearing impairment |
| 61 | Hypertonia |
| 62 | Hypotonia |
| 63 | Intellectual disability |
| 64 | Joint hypermobility |
| 65 | Joint laxity |
| 66 | Laryngeal web |
| 67 | Macrocephaly |
| 68 | Microcephaly |
| 69 | Motor delay |
| 70 | Muscle weakness |
| 71 | Poor growth |
| 72 | Recurrent otitis media |
| 73 | Scoliosis |
| 74 | Seizure |
| 75 | Short attention span |
| 76 | Sleep disturbance |
| 77 | Stroke |
| 78 | Tremor |
| 79 | Visual impairment |

| N | Clinical Phenotypes combined 1 |
|---|---|
| 1 | Abnormal vertebral morphology |
| 2 | Abnormality of temperature regulation |
| 3 | Abnormality of the mitochondrion |
| 4 | Anxiety |
| 5 | Ataxia |
| 6 | Behavioral abnormalities |
| 7 | Delayed speech and language development |
| 8 | Depression |
| 9 | Developmental regression |
| 10 | EEG abnormality |
| 11 | Encephalopathy |
| 12 | Feeding difficulties |
| 13 | Global developmental delay |
| 14 | Intellectual disability |
| 15 | Laryngeal web |
| 16 | Scoliosis |
| 17 | Seizure |
| 18 | Short attention span |
| 19 | Sleep disturbance |
| 20 | Stroke |
| 21 | Tremor |
| 22 | abnormality of body parts ( ear, jaw, inner ear, hairline, finger, dentition) |
| 23 | heart function related (EKG, cardiovescular) |
| 24 | Motor function |
| 25 | Abnormality of liver |
| 26 | Skin problems |
| 27 | Inner organs (lung) |
| 28 | Brain/nervous system |
| 29 | Vision related |
| 30 | Muscle related |
| 31 | Autism related |
| 32 | Movement? |
| 33 | Joint? |
| 34 | Tonia? |
| 35 | Hearing problem |
| 36 | Growth problems |
| 37 | Cephaly probems |

| N | Clinical Phenotypes combined 2 |
|---|---|
| 1 | Abnormal vertebral morphology |
| 2 | Abnormality of temperature regulation |
| 3 | Abnormality of the mitochondrion |
| 4 | Anxiety, Short attention span, Depression |
| 5 | Ataxia |
| 6 | Behavioral abnormalities |
| 7 | Developmental regression |
| 8 | Feeding difficulties |
| 9 | Intellectual disability |
| 10 | Scoliosis and Joint |
| 11 | EEG abnormality, Seizure and Encephalopathy |
| 12 | Sleep disturbance |
| 13 | Stroke |
| 14 | Motor function, Movement and Tremor |
| 15 | Neurodevelopmental (Delayed speech and language development, Global developmental delay) |
| 16 | Sensory (Vision related, Hearing problem) |
| 17 | Abnormality of peripheral body parts ( ear, jaw, inner ear, hairline, finger, dentition, skin) |
| 18 | Heart function related (EKG, cardiovescular) |
| 19 | Inner organs (lung, liver, laryngeal web) |
| 20 | Brain/nervous system |
| 21 | Muscle related and Tonia |
| 22 | Autism related |
| 23 | Growth problems |
| 24 | Cephaly probems |

| N | Clinical Phenotypes final |
|---|---|
| 1 | Ataxia |
| 2 | Hypotonia |
| 3 | Tremor |
| 4 | Seizure + EEG abnormality |
| 5 | Delayed language |
| 6 | DDID |
| 7 | Motor |
| 8 | Encephalopathy |
| 9 | Sensory problem |
| 10 | Abnormality of face |
| 11 | Joint |
| 12 | Mood/attention/psych |
| 13 | Brain atrophy |
| 14 | Conjugate eye movement |
| 15 | Motor delay |
| 16 | Movement |
| 17 | Muscle |
| 18 | Gait |
