## Supplementary figures and images for "Characterization of the functional and clinical impacts of CACNA1A missense variants found in neurodevelopmental disorders"

### Supp. Fig. 1

#### Slide 1
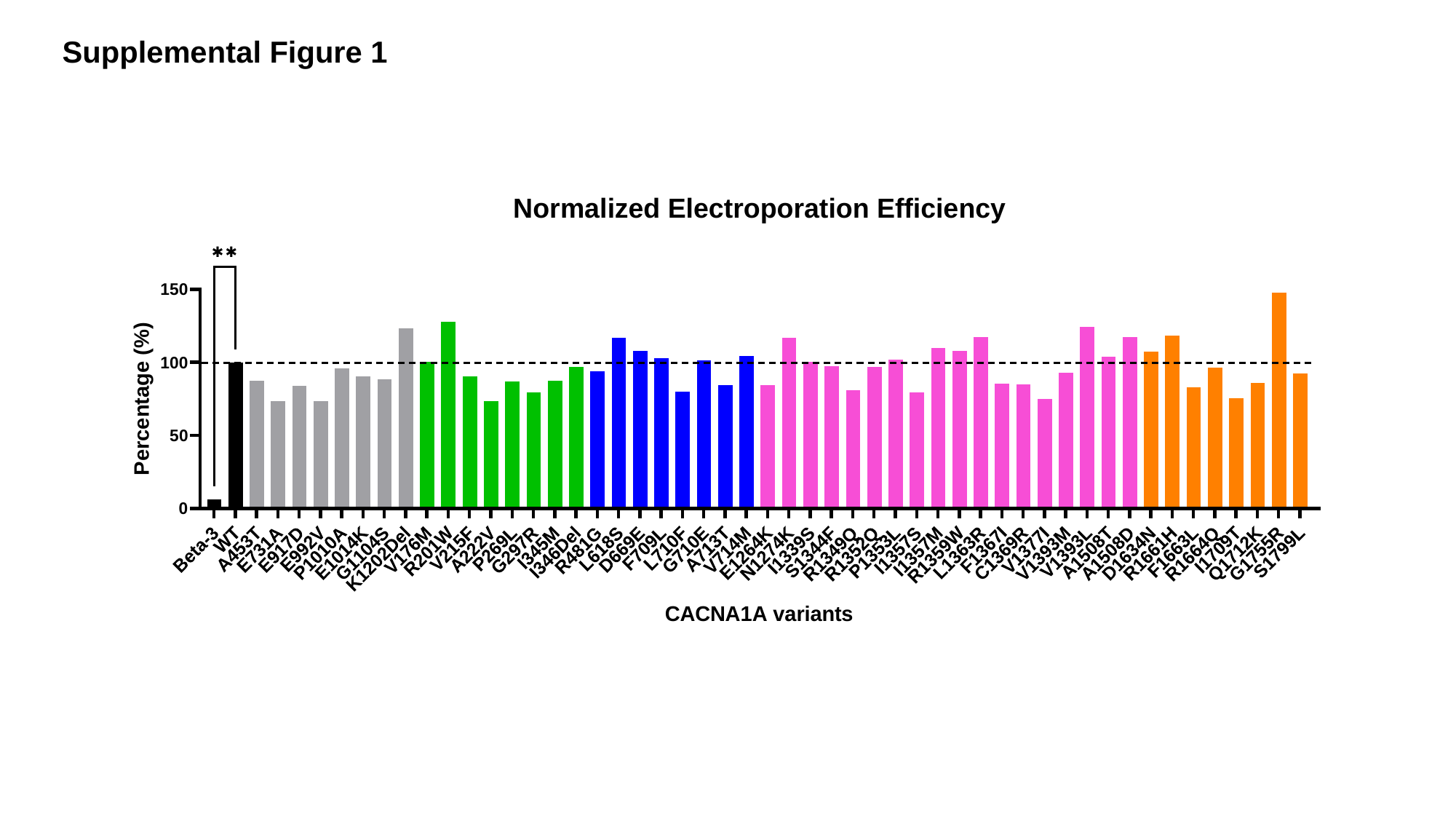

Supplemental Figure 1

### Supp. Fig. 2

#### Slide 1
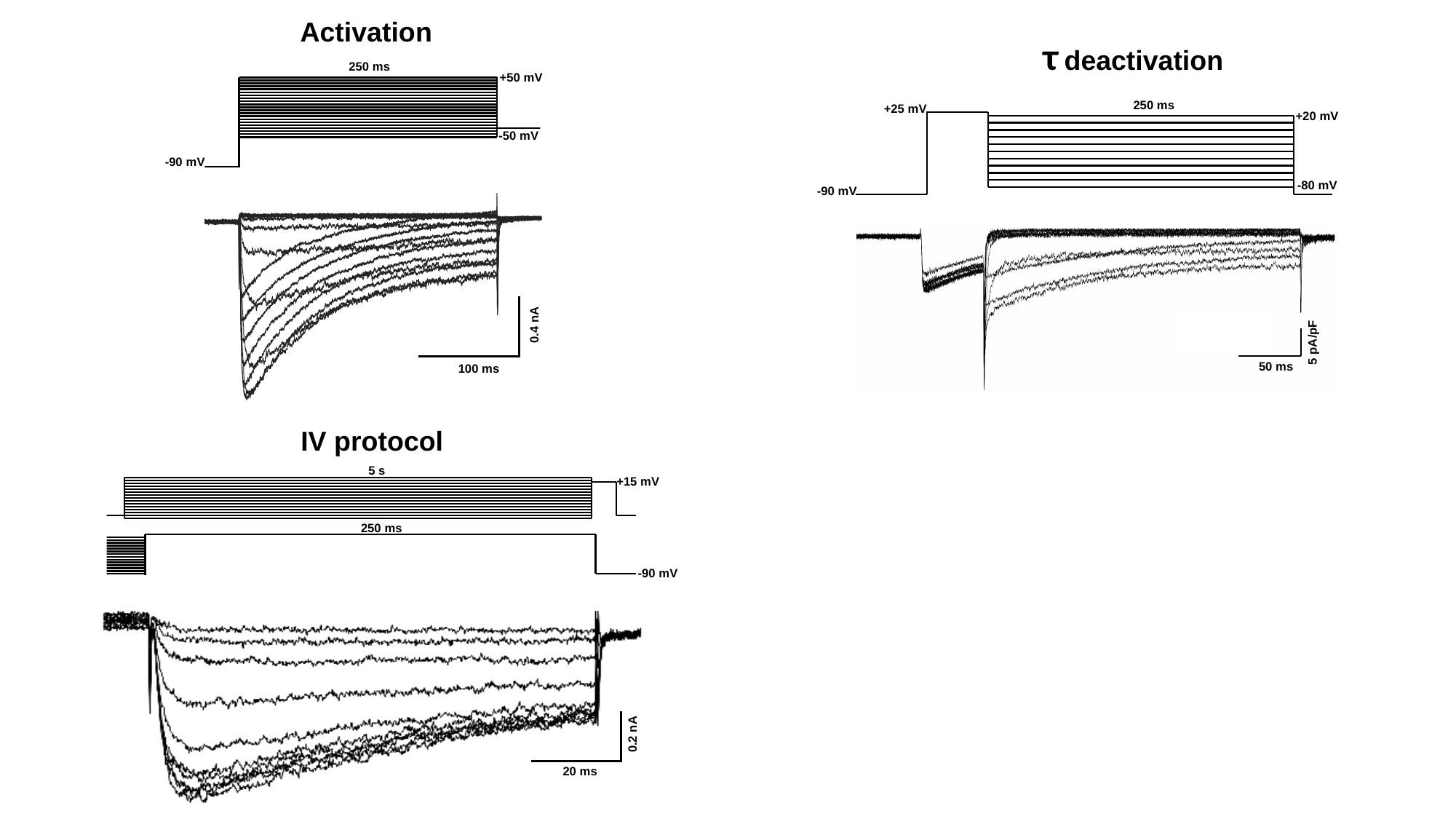

Activation
τ deactivation
250 ms
+50 mV
-50 mV
-90 mV
0.4 nA
100 ms
250 ms
+25 mV
+20 mV
-80 mV
-90 mV
5 pA/pF
50 ms
IV protocol
5 s
+15 mV
250 ms
-90 mV
0.2 nA
20 ms

### Supp. Fig. 3

#### Slide 1
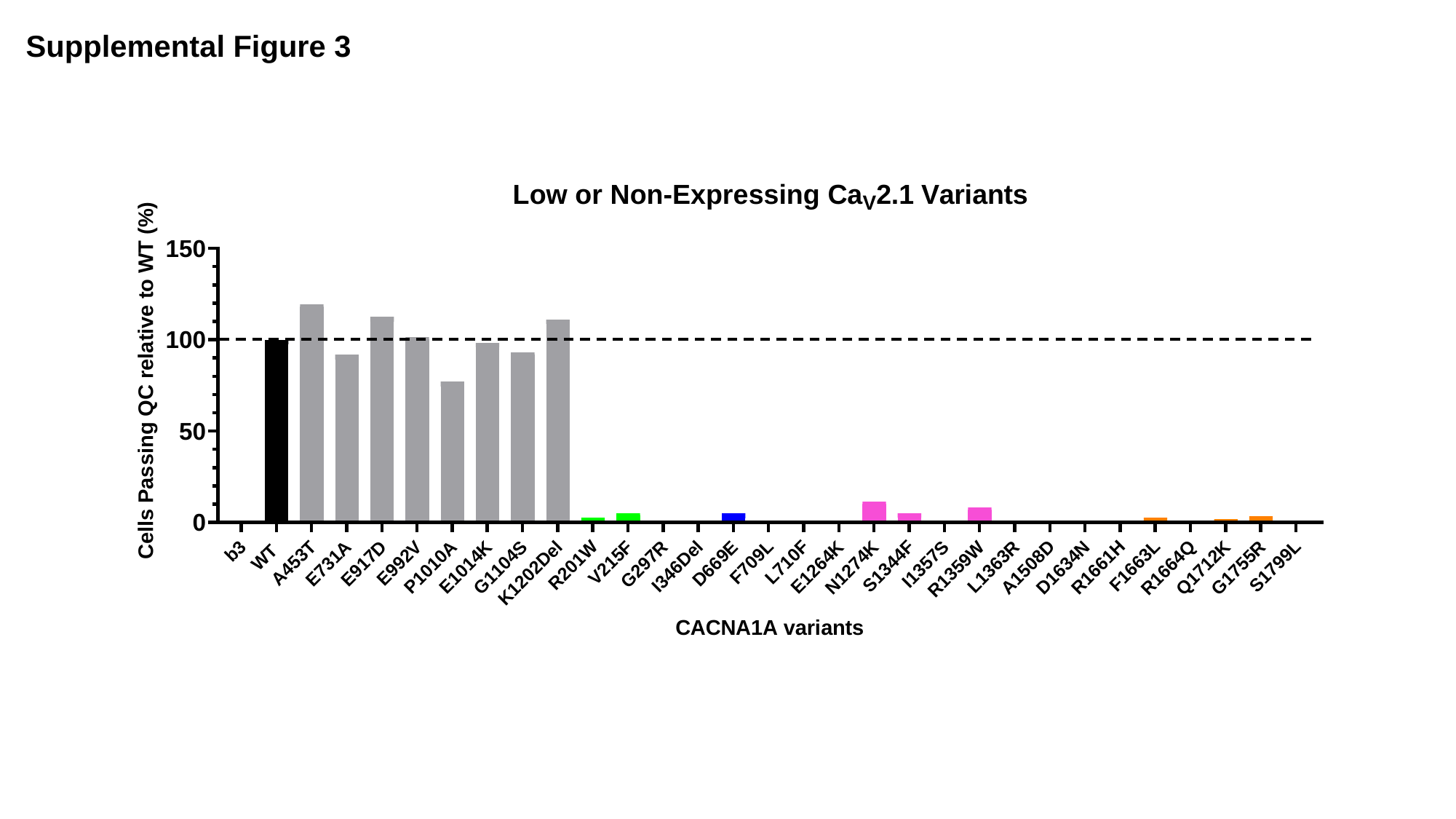

Supplemental Figure 3

### Supp. Fig. 4

#### Slide 1
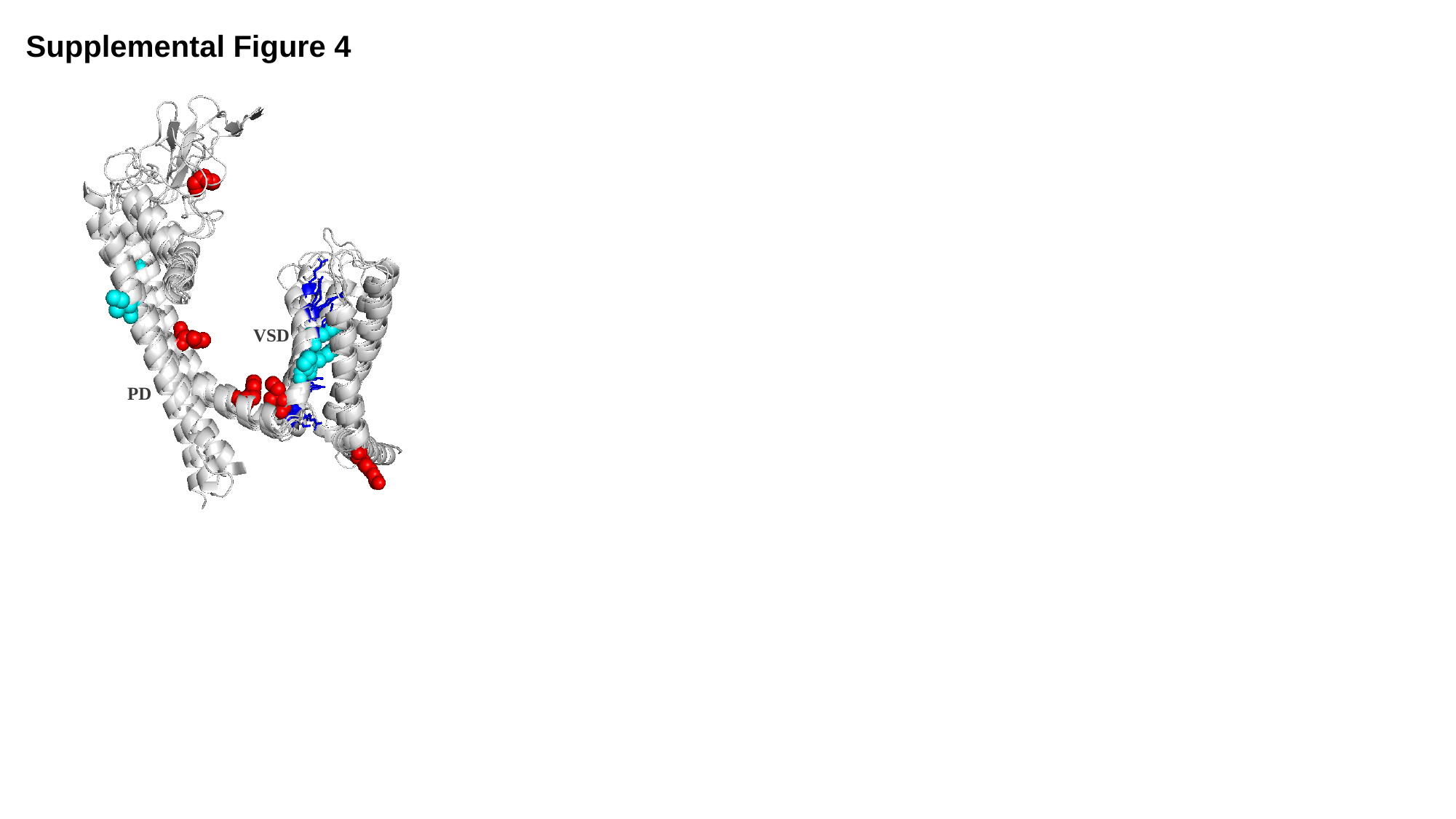

Supplemental Figure 4
VSD
PD
