## Supplementary material for "Characterization of the functional and clinical impacts of CACNA1A missense variants found in neurodevelopmental disorders": Supp. Fig. 5

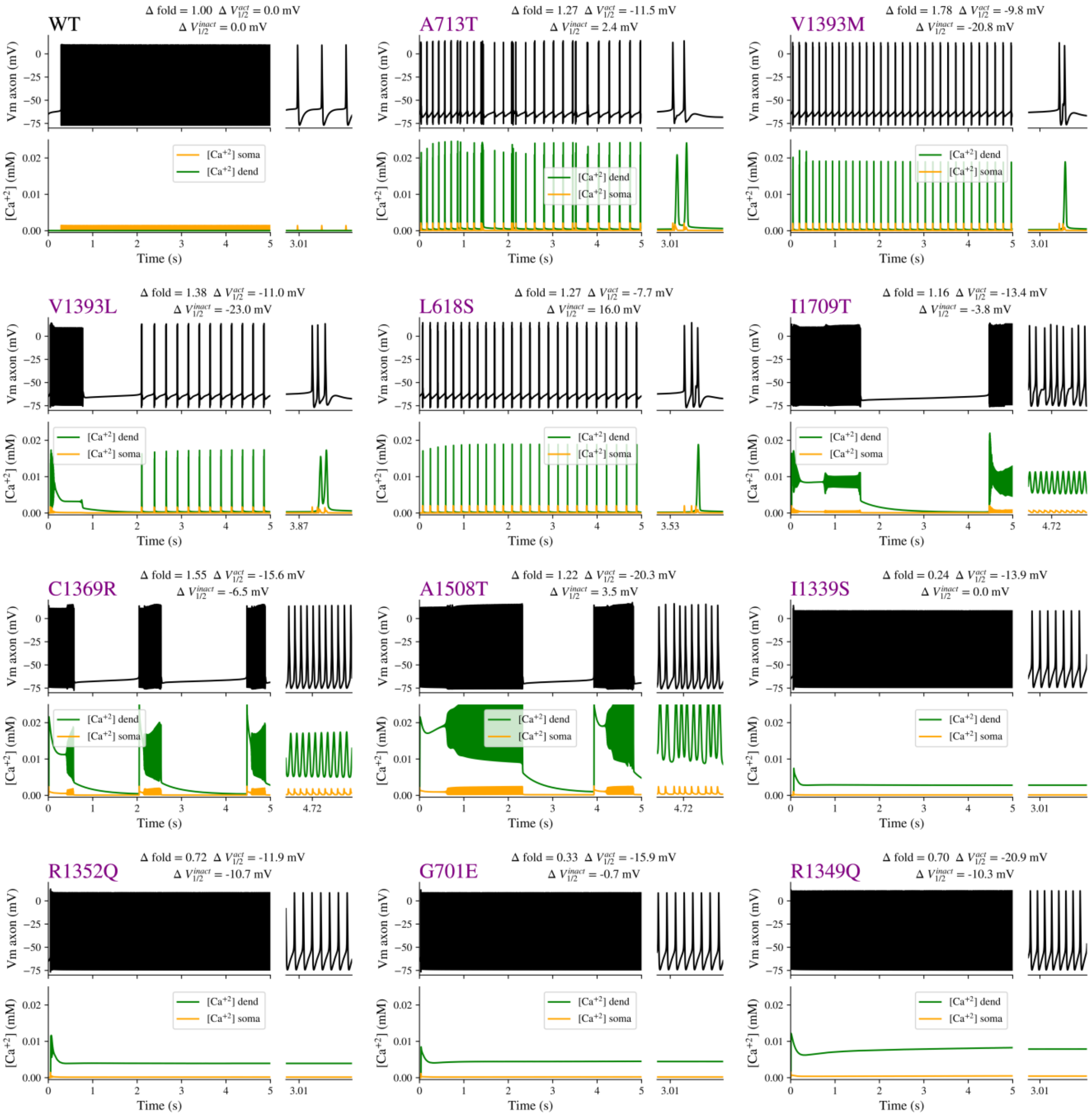


**Figure S5. Sample of simulated axonal voltage traces and intracellular $Ca^{2+}$ concentration changes for all the variants that presented complex firing patterns in Fig. 4 (fuchsia dots).** In the upper panel are represented the voltage changes vs time during the 5-second simulations (black trace). In the lower panel the intracellular calcium concentration in the dendrites (green trace) and soma (yellow trace) are plotted at the same scale as the voltage. Voltage and $Ca^{2+}$ plots are followed by a 50 ms inset at the time points specified in the X-axis to appreciate better faster transitions for all the recordings. At the top of each panel are listed the details how the Ca2.1 conductance was modified during the simulation to represent the changes associated with each variant. Δfold for fraction of total conductance (WT value 1.0), $\Delta_{ACT\text{-}V1/2}$ for shift in the voltage activation midpoint (WT = 0 mV), and $\Delta_{INACT\text{-}V1/2}$ for changes in voltage inactivation midpoint (WT = 0 mV).
